# Senescence rewires microenvironment sensing to facilitate anti-tumor immunity

**DOI:** 10.1101/2022.06.04.494739

**Authors:** Hsuan-An Chen, Yu-Jui Ho, Riccardo Mezzadra, Jose M. Adrover, Ryan Smolkin, Changyu Zhu, Wei Luan, Alexandra Wuest, Sha Tian, Xiang Li, Ronald C. Hendrickson, Mikala Egeblad, Direna Alonso-Curbelo, Scott W. Lowe

## Abstract

Cellular senescence involves a stable cell cycle arrest coupled to a secretory program that, in some instances, stimulates the immune clearance of senescent cells. Using an immune competent tumor model in which senescence triggers CD8 T cell-mediated tumor rejection, we show that senescence also remodels cell surface proteome to alter how they sense environmental factors, as exemplified by Type II interferon gamma (IFN-γ). Compared to proliferating cells, senescent cells upregulate IFN-γ receptor, become hypersensitized to microenvironmental IFN-γ, and more robustly induce antigen presenting machinery -effects also recapitulated in human tumor cells treated with senescence-inducing drugs. Disruption of the IFN-γ sensing by senescent cells blunts their immune-mediated clearance without disabling their characteristic secretory program or immune cell recruitment. Our results demonstrate that senescent cells have an enhanced ability to both send and receive environmental signals, and imply that each process is required for their effective immune surveillance.

**SIGNIFICANCE:** Our work identifies a novel interplay between tissue remodeling and tissue sensing programs that can be engaged by senescence in advanced cancers to render tumor cells more visible to the adaptive immune system. This new facet of senescence establishes reciprocal heterotypic signaling interactions that can be induced therapeutically to enhance anti-tumor immunity.

## INTRODUCTION

Cellular senescence is a stress response program characterized by a stable cell cycle arrest and a secretory program capable of remodeling the tissue environment (1). In normal tissues, senescence contributes to tissue homeostasis during wound healing; however, in aged or damaged tissues, the aberrant accumulation of senescent cells can cause chronic inflammation and reduced tissue regenerative capacity (2–4). In cancer, senescence has been shown to mediate both beneficial and detrimental effects on tissue biology. In certain contexts, senescence provides a barrier to oncogene-initiated tumorigenesis and may contribute to the anti-tumor activity of a range of cancer therapies (5, 6). Conversely, the aberrant accumulation of senescent cells following such therapies can stimulate tumor resistance, tumor progression and metastasis (7, 8). Therefore, senescence appears to have both beneficial and detrimental effects on tissue biology through effects that remain poorly understood.

One facet of the senescence program that is likely to contribute to such diverse biology is the senescence-associated secretory phenotype [SASP, ref: (9)]. SASP is activated through a global chromatin remodeling process that evolves over time and is controlled by key epigenetic regulators such as Brd4 and pro-inflammatory transcription factors such as NF-κb and C/EBP-β (10–12). This, in turn, leads to the induction of genes that encode tissue remodeling proteins such as matrix metalloproteinases, growth factors, and fibrolytic factors known to play crucial roles in wound healing process (3,13,14). Other SASP components include a diverse set of chemokines and cytokines that can alter the composition and state of immune cells within the tissue, sometimes leading to the immune-mediated targeting and clearance of the senescent cells themselves (15, 16). Nonetheless, the aberrant accumulation of senescent cells in many pathological contexts implies that immune mediated clearance is not a universal outcome of SASP and raises the possibility that additional mechanisms dictate the paradoxically beneficial and detrimental effects of senescence in tissue biology and immune surveillance (17–19).

Certainly, senescence-associated immune surveillance has potent anti-cancer effects, though the precise mechanisms depend on context. For example, cells undergoing oncogene induced senescence can be eliminated by NK cells or the combined action of CD4+ T cells and macrophages eliminates premalignant liver cells (10, 16). Moreover, while most tumor cells evade senescence during tumor evolution, the process can also be reengaged in advanced disease stages by treatment with certain cytotoxic or targeted cancer therapies, reactivating innate immuntiy or re-sensitizing tumors to the activity of CD8-positive T cells and/or immune checkpoint blockade in some settings (20–22). Still, in other contexts, therapeutic induction of senescence and SASP does not lead to immune surveillance but, in stark contrast, stimulates inflammation to favor tumor progression and relapse (8,19,23). Therefore, understanding the mechanisms by which senescent tumor cells become visible to the immune system may facilitate strategies to elicit anti-tumor immunity in patients.

Here, we set out to establish principles that modulate the immune recognition and clearance of senescent cells to identify actionable senescence mechanisms that may be exploited to improve the immune control of cancer. To this end, we developed a novel ‘senescence-inducible’ model in which liver cancer cells can be selectively switched to a senescent state through genetic modulation of endogenous p53, thereby avoiding the confounding effects of senescence-inducing therapies on immune cells or other components of the tissue environment. Using this system, we reveal that, in addition to the SASP, senescence drives a major remodeling of the cell surface proteome in a manner predicted to fundamentally alter the way cells sense and respond to environmental signals, exemplified herein through a hypersensitivity to microenvironmental type II IFN (IFN-γ). This process enables a robust upregulation of the antigen processing and presenting machinery in tumor cells that renders them more susceptible to CD8 T-cell mediated killing. Thus, our results uncover a rewired tissue sensing program in senescent cells that acts in concert with SASP to establish heterotypic interactions with their tissue environment that boost their immunogenic potential to facilitate immune-mediated clearance.

## RESULTS

### A p53-restorable immunocompetent tumor model to study senescence surveillance

To study how senescence reprograms cellular and tissue states, we exploited the hydrodynamic tail vein injection (HTVI) technique (24) to generate a senescence-inducible liver cancer model controlled by a tumor-specific, restorable p53 short-hairpin RNA (shRNA). Specifically, adult liver hepatocytes of immunocompetent Bl/6 mice were transfected *in vivo* with a sleeping beauty SB13 transposase vector and two transposon constructs (encoding <u>N</u>rasG12D-IRES-rtTA and TRE-tRFP-<u>s</u>h<u>p</u>53, or “NSP”) which integrate in the genome. In this Tet-On system, endogenous p53 is suppressed in the presence of doxycycline (Dox) through the activation of inducible shRNA linked to RFP (**Fig. 1A**). Consistent with the co-occurrence of mutations that inactivate *TP53* and activate cell proliferation signaling pathways (e.g. PI3K/AKT and RAS/MAPK cascades) in human liver tumors, the cooperation between oncogenic RAS and suppression of p53 led to hepatocyte transformation, with most mice developing tumors with poorly differentiated features 5-8 weeks after HTVI. Transcriptional profiling using RNA-seq revealed that these tumors resemble the ‘proliferation class’ of human hepatocellular carcinoma [HCC, ref. (25)], characterized by their enriched frequency of *TP53* mutations and worst prognosis [Supplementary Fig. S1; ref. (26, 27)].

**Figure 1.**
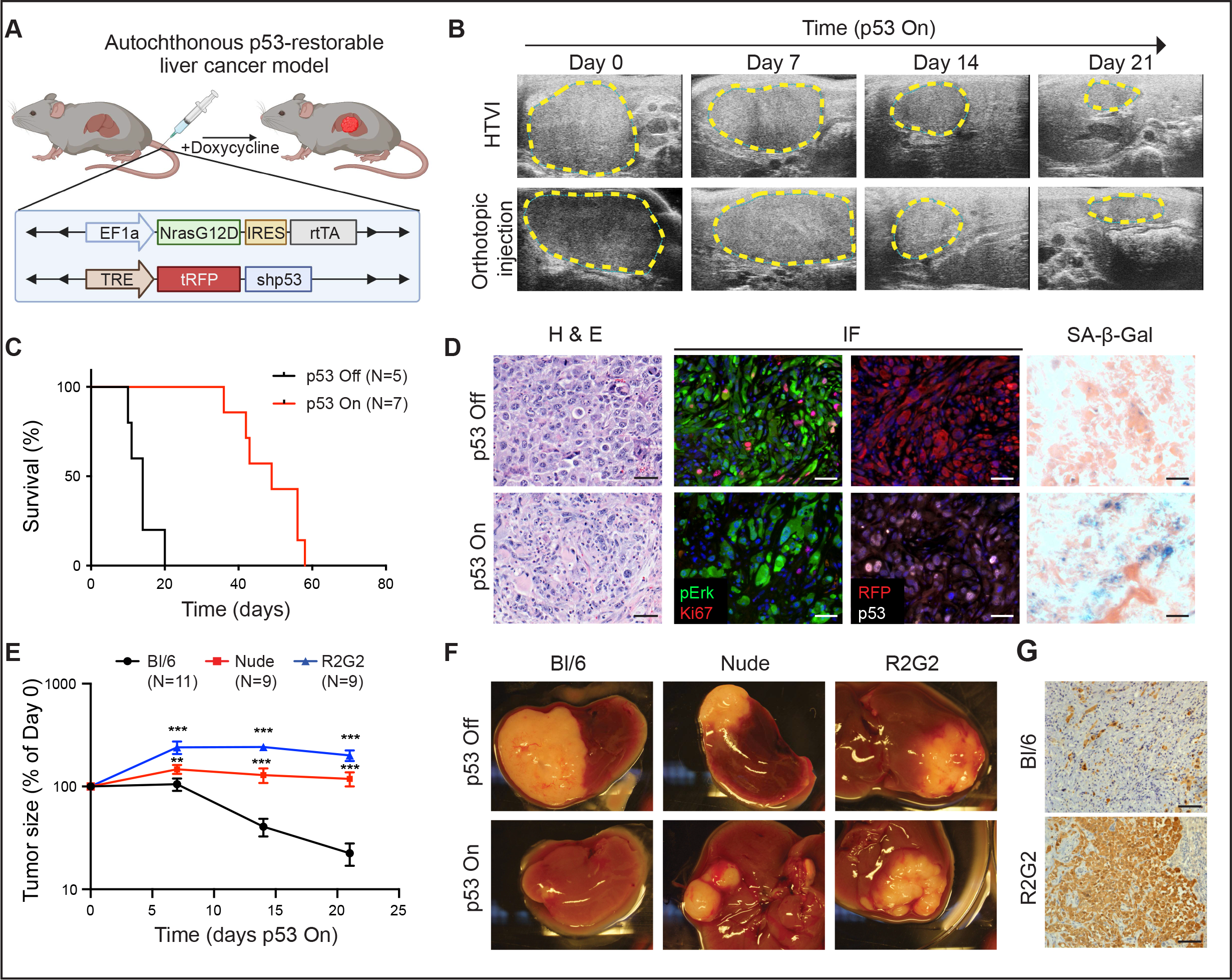
A p53-restorable tumor model to study senescence immune surveillance. A, Generation of the p53-restoratable, NRAS-driven mouse liver cancer model using the sleeping beauty transposon system delivered through hydrodynamic tail vein injection (HTVI). B, Representative ultrasonogram of HTVI and orthotopic injection liver cancer models at indicated time after p53 restoration. C, Survival analysis of mice in the HTVI model. D, Representative haematoxylin and eosin (H&E), immunofluorescence (IF) and senescence-associated β-Gal (SA-β-Gal) staining of p53-suppressed (p53 Off) and -restored (p53 On for 14 days) tumor sections generated from the HTVI model. Scale bar, 50 µm. E to G, Orthotopic injection of GFP-luciferase vector-transduced NSP tumor cells into the livers of immunocompetent and –deficient mouse strains. E, Tumor size change measured by ultrasound upon p53 restoration. R2G2, Rag2-Il2rg double knockout mouse. Data are presented as mean ± s.e.m. N ≥ 9 for each strain. F, Representative macroscopic pictures at 21 days of p53 On or end-point p53 Off tumor. G, Representative immunohistochemistry (IHC) staining of GFP-labeled tumor cells at day 21 upon p53 restoration. Scale bar, 100 µm.

Based on previous work (15), we anticipated that reactivation of p53 in the above *in vivo* system would switch tumor cells to a senescent state and engage tumor-suppressive host immunity mechanisms. Accordingly, dox withdrawal triggered dramatic tumor regressions over the course of several weeks, leading to prolonged animal survival (**Fig. 1B and 1C**). Analysis of the tumors at 14 days post dox withdrawal revealed the expected downregulation of the p53-shRNA (as visualized by the linked RFP reporter) and accumulation of senescence-associated-β-galactosidase (SA-β-gal) without any notable effects on the RAS-effector p-ERK (**Fig. 1D**). Similarly, SA-β-gal positivity together with a concomitant reduction in proliferation and increase in the p53 target p21 was noted in tumor cells explanted in culture (Supplementary Fig. S2A-S2D). Importantly, injection of such cultures (maintained on Dox to keep p53 off) into immunocompetent mice (on a Dox diet) produced synchronous and focal tumors within three weeks that regressed with similar kinetics as the primary tumors upon Dox withdrawal (**Fig. 1B**; Supplementary Fig. S2E-S2I). Of note, control experiments using a Tet-Off system or expressing a constitutive p53 shRNA ruled out the possibility that Dox itself had any effect on tumor behavior in our model (Supplementary Fig. S2J and S2K). Therefore, this system allows for the efficient induction of senescence in tumor cells without resorting to therapies that can also alter the host. Given its added flexibility, we used the orthotopic transplant model (hereafter referred to as ‘NSP’) for many of the mechanistic studies described below.

As anticipated, the marked tumor regression noted above were immune mediated. Hence, NSP tumors that arose following transplantation into immunocompromised Nude and *Rag2^-/-^Il2rg^-/-^*(R2G2) mice underwent a prominent cytostatic response but failed to regress, with R2G2 animals showing the most profound effects (**Fig. 1E-1G**; Supplementary Fig. S2L and S2M). As Nude mice are defective in adaptive immunity whereas R2G2 are also compromised for aspects of innate immunity, these results imply that the adaptive immune system is essential for efficient tumor regression in the model and establish a well-controlled experimental context to explore the mechanistic basis for these effects.

### Senescence triggers an immune evasion-to-immune recognition tumor switch

To characterize the tumor suppressive paracrine effects of senescence, we next characterized the immune microenvironments of tumors harboring p53-suppressed (referred to as “proliferating’) and p53-restored (referred to as “senescent”) tumor cells. Preceding tumor regressions, lesions harboring senescent tumor cells showed a ∼1.8-fold increase in total CD45+ immune cells compared to proliferating controls [**Fig. 2A**; refs. (15, 16)] that, upon immunophenotyping, involved a prominent proportional increase in lymphocytes (B cells, CD4+ and CD8+ T cells) and decrease of the Gr-1+ myeloid-derived suppressor cells/neutrophils (CD11b+Gr1+Ly6C low) (**Fig. 2B**; Supplementary Fig. S3A). Among the T cell population, accumulating CD8 T cells showed markers of antigen experience (CD69+, CD44+, PD1+) and harbored an increased population of effector cells (CD44+CD62L-) [**Fig. 2C**; refs. (28, 29)]. Overall remodeling of immune cell infiltrates led to a significant increase in the CD3:neutrophil ratio for tumors harboring senescent cells (Supplementary Fig. S3B). This profound effect is consistent with similar increases in the CD3:neutrophil ratio which have been associated with immune reactivity in human liver tumors (30) and could be clearly visualized using three dimension (3D) imaging after tissue clearing [**Fig. 2D**; Supplementary Fig. S3C and S3D; Supplementary video S1; ref. (31)].

**Figure 2.**
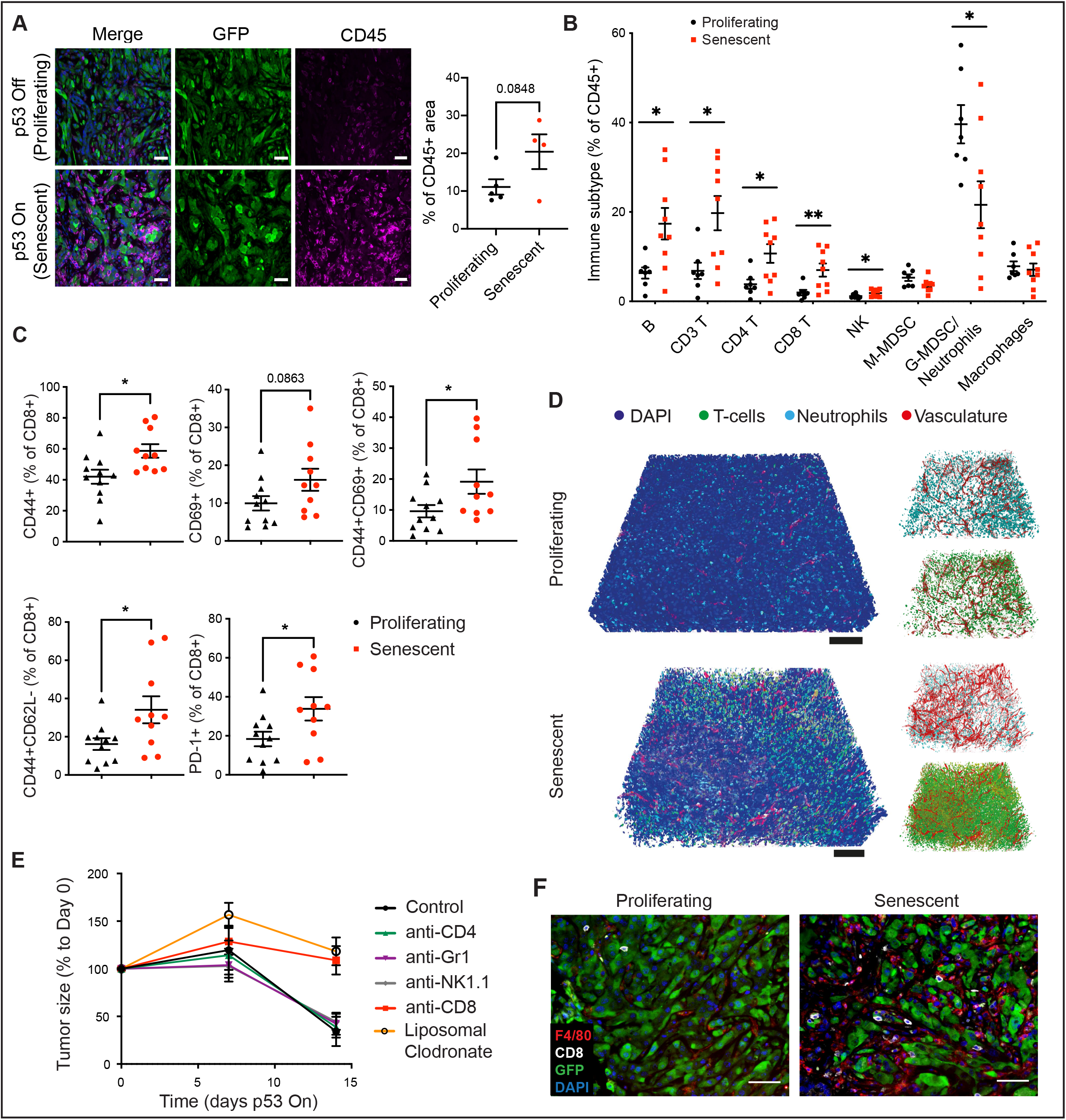
Senescence triggers an immune evasion-to-immune recognition tumor switch. A, Representative images of CD45 and GFP staining marking immune cells and tumor cells, respectively in p53-suppressed and p53-restored tumor (7 days after p53 restoration). Right panel is the quantification of the area of CD45+ staining calculated from 3 random fields per mouse. Each dot represents a mouse. B, Flow cytometry analysis of global immune landscape in orthoptopic NSP liver tumor model. Immunophenotyping of senescent tumors is performed 9 days after Dox withdrawal, a time point when the senescent state is fully established yet, preceding the tumor regression. Data is pooled from 2 independent experiments with n=7 in the proliferating group and n=9 in the senescent group. C, Flow cytometry analysis of CD8 T cells. Data is pooled from 2 independent experiments with n=11 in the proliferating and n=10 in the senescent groups. D, Representative tissue clearing images of the orthotopic NSP liver tumors. T cells, neutrophils and vasculature are labeled by CD3, MPO and CD31 staining, respectively. E, Tumor size change measured by ultrasound upon p53 restoration in mice after depleting specific immune cell types. F, Representative immunofluorescence images of CD8 T cells and F4/80 positive macrophages staining in the orthotopic NSP liver tumor. Data is presented as mean ± s.e.m. Scale bar, 100 µm. A two-tailed student t-test is used. *p < 0.05; **p < 0.01.

To pinpoint the specific immune cell types responsible for immune surveillance of senescent tumor cells, we generated parallel cohorts of mice harboring orthotopic NSP tumors and performed Dox withdrawal along with simultaneous treatment with immune cell-depleting agents, monitoring tumor regression over time. Whereas blocking antibodies targeting neutrophils/monocytes (Gr1), NK cells (NK1.1), and CD4 positive T cells (GK1.5) had no effect, depletion of CD8 T cells (2.43) and macrophages (using liposomal clodronate which selectively targets macrophages (CD11b+F4/80+) but not cDC (CD11b-CD11c+MHC-II+CD103+); ref. ((32, 33))) markedly impaired tumor regression (**Fig. 2E; Supplementary Fig. S3E**). Interestingly, histological analyses revealed CD8+ T cells and F4/80+ macrophages were frequently co-enriched following senescence induction in both primary HTVI and orthotopic transplant tumors (**Fig. 2F**; Supplementary Fig. S3F and S3G), suggesting a CD8-dependent immune response involving cooperativity with macrophages clears p53-reactivated senescent tumor cells. Thus, p53 reactivation induces tumor cell senescence leading to an abrupt switch from immune evasion to immune surveillance, productive anti-tumor immunity and, ultimately, tumor rejection.

### Senescence remodels tissue sensing programs and cell-surfaceome landscape

We next set out to exploit our model understand the molecular mechanisms responsible for rendering senescent tumor cells visible to the immune system. Previous studies demonstrate that senescence induction involves a chromatin remodeling program that silences proliferative genes and activates many genes encoding for SASP factors, with the latter program being largely dependent on the enhancer reader, BRD4 (10). We therefore performed transcriptional profiling experiments on NSP cells under proliferating (p53-suppressed) versus senescent (p53-restored) conditions in the absence and presence of JQ1, a drug that inhibits BRD4 function (Supplementary table S1). Consistent with expectations, p53 restoration dramatically reduced the expression of proliferative genes and induced the expression of well-known SASP factors [**Fig. 3A**; Supplementary Fig. S4A and S4B; ref: (7)], including several cytokines known to stimulate T cells (*Cxcl16, Il18*) or macrophage activation and recruitment (*Csf2*, encoding protein GM-CSF) or previously linked to senescence (*Igfbp7, Igfbp3, Pdgfa*). As anticipated from previous work (10), many of the upregulated SASP transcripts (∼65%) were BRD4-dependent (Supplementary Fig. S4B). Similarly, a range of growth factors and immune modulators were secreted from the senescent cells as assessed by multiplexed cytokine assays, including the T cell and macrophage attractants CCL5, CXCL9, and GM-CSF as well as vasculature remodeling factor VEGF (Supplementary Fig. S4C). Therefore, the induction of senescence in p53-restored NSP tumor cells is associated with a robust SASP, consistent with marked remodeling of the tumor immune ecosystem characteized above.

**Figure 3.**
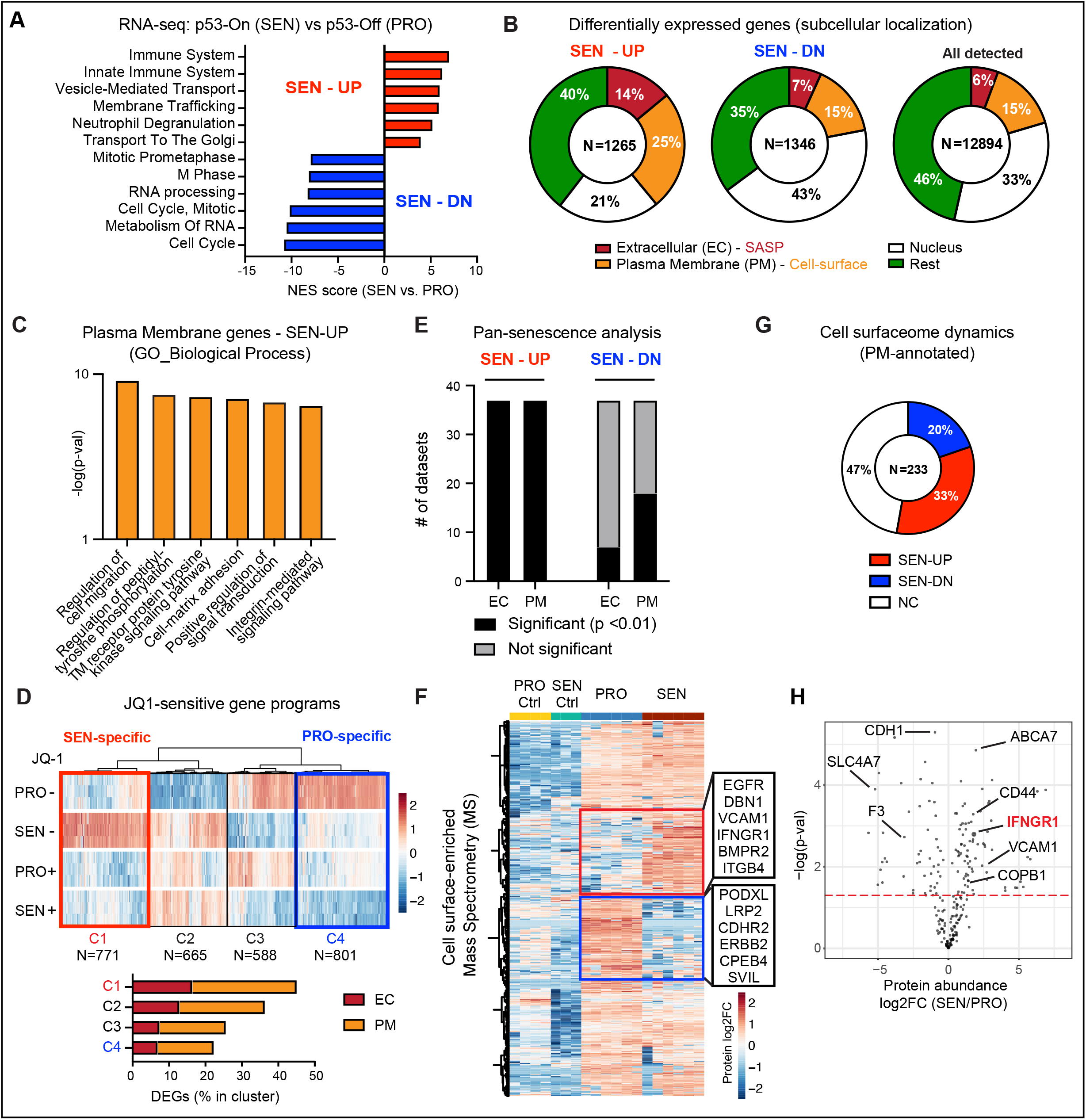
Senescence remodels tissue sensing programs and cell-surfaceome landscape. A, GSEA (Reactome) of RNA-Seq data from proliferating (PRO, p53 Off) vs. senescent (SEN, p53 On for 8 days) NSP liver tumor cells in vitro. B, Subcellular localization of DEGs (p < 0.05; fold change > 2) all detected genes (TPM > 1) from RNA-seq. C, Gene Ontology (GO) analysis of DEGs encoding plasma membrane proteins upregulated in senescent cells. D, Transcriptomic analysis of all differential expressed genes (DEGs, proliferating vs. senescent) in the presence or absence JQ1 treatment. C1 cluster (in red) contains the senescence-specific genes sensitive to JQ1 and C4 cluster (in blue) contains the proliferation-specifiic genes sensitive to JQ1. E, Meta-analysis of RNA-seq datasets from SENESCOPEDIA by performing subcellular localization of DEGs (same as Fig. 2D) and Fisher exact test to examine the relative enrichment of up- and downregulated EC/PM-DEGs deviated from the random distribution. See also supplementary Fig. S4E and S4F. F, Mass spectrometry (MS) analysis of plasma membrane-enriched proteome in proliferating and senescent cells. Protein level is normalized to mean expression of the protein of all samples. Controls are the samples without biotin labeling serving as background. Red and blue boxes represent proteins enriched in senescent and proliferating cells respectively. N=6 for both senescent and proliferating experimental group, and N=3 and 4 respectively for their control. G, Distribution of up- and downregulated Genecard-annotated plasma membrane (PM) proteins profiled by MS. H, Volcano plot of Genecard-annotated plasma membrane proteins profiled by MS. EC, extracellular; PM, plasma membrane.

Strikingly, further examination of the subcellular localization for differentially expressed genes (DEGs) revealed that senescent tumor cells not only increased their expression of secreted (‘extracellular’, EC) SASP factors, but also displayed major changes in the expression levels of transcripts encoding surface proteins (‘plasma membrane’, PM) (**Fig. 3B**). Indeed, 25% of total upregulated DEGs encoded for PM proteins, a significant enrichment that deviated from the random distribution (15%) (**Fig. 3B**). Dynamic PM-DEGs were linked to protein tyrosine kinase signaling transduction (*Nrp1, Egfr*), cytokine receptor activity (*Ifngr1*), ECM receptors (*Itgb3*, *Cd44*) and ion transporters (*Slc12a1*, *Slc24a3*) and captured known senescence-associated molecules (*Cd44, Vcam1, and Itgb3*), suggesting senescent cells may have a enhanced capability to interact with and sense their environment [**Fig. 3C**; Supplementary Fig. S4D; refs: (34–36)].

Interestingly, the senescence-associated increase in the expression of many of these PM proteins was blunted by JQ1, suggesting that their induction may be part of the broader chromatin remodeling program coupled to SASP (**Fig. 3D**). Of note, profound changes in transcription of genes encoding PM proteins also occurred in p53-deficient NSP tumor cells treated with the senescence-inducing drug combination trametinib and palbociclib [Supplementary Fig. S4E; Supplementary table S1; ref: (20)] and in a series of 13 genetically-diverse human cancer lines induced to senesce by various triggers (SENESCOPEDIA) (37), indicating that the remodeling of PM factors may be a universal feature of senescent cells, and not a phenomenaon exclusively linked to our p53-reactivation system (**Fig. 3E**). This was particularly robust for upregulated (but not downregulated) PM-DEGs, reminiscent to effects observed for extracellular (EC) SASP factors (**Fig. 3E**; Supplementary Fig. S4F). Therefore, the markedly altered expression of cell surface proteins we observed in our model extends beyond p53-induced senescence and may be a hallmark of the senescent state.

To validate the global remodeing of PM factors in senescence at the protein level, we performed surface proteomics on isogenic proliferating and senescent NSP tumor cells, using a biotin-labeling enrichment method, where cell surface proteins were labeled with membrane-impermeable biotin, purified, and subjected to mass spectrometry (38) (**Fig. 3F**; Supplementary Fig. S4G). The method was robust: results indicated a strong correlation between biological replicates under each condition (Supplementary Fig. S4H), with detected proteins being enriched for annotated plasma membrane proteins by 60% after induction of p53-induced senescence. Of 887 proteins that were reproducibly detected, > 50 % were differentially expressed. While many of these differentially expressed proteins correlated well with the directionality observed in our transcriptional profiling data, a subset showed no significant change in transcript levels (Supplementary Fig. S4I). Annotated cell surface proteins detected by mass spectrometry upon senescence induction included several previously linked to senescence (e.g. CD44, VCAM1), different growth factor and cytokine receptors (e.g. EGFR, ICAM1 and IFNGR1), and other less characterized factors (**Fig. 3G and 3H**; Supplementary Fig. S4J and S4K). Of note, the set of cell surface-enriched proteins identified in our model showed limited overlap with those identified in human fibroblasts undergoing oncogene-induced senescence (39), suggesting heterogeneity between cell types or senescence triggers. Regardless, these results show that in addition to a rewiring in their secretory program, senescent cells undergo profound changes in the content and abundance of cell surface proteins, and imply that senescent cells acquire distinctive microenvironment-sensing traits that may influence their state and fate in vivo.

### Senescent cells are primed to sense and amplify IFN-γ signaling

To identify pathways distinctly-altered in senescent cells that could, in principle, influence how senescent cells interact with tumor microenvironment (TME) cells. we next mined transcriptional and proteomic datasets for senescence-associated changes that may be potentially involved in mediating the rboust immune-mediated elimination phenotype observed *in vivo*. Interestingly, we noted that among the top 5 annotated pathways that were both enriched during senescence and dependent on cell state-specific enhancer programs (i.e. JQ1-sensitive) were factors linked to type II interferon (IFN) signaling, a cascade crucial for provoking anti-tumor immunity (40) (Supplementary Fig. S5A). Specficially, type II IFN-gamma (IFN-γ) binding to the IFNGR (a heterodimer of the IFNGR1 and IFNGR2 subunits) signals through the JAK-STAT pathway leading to phosphorylation of the transcriptional regulator STAT1 which, upon entry into the nucleus, binds to interferon-<u>g</u>amma <u>a</u>ctivated <u>s</u>ites (GAS) to drive the expression of interferon stimulated genes (ISGs). This drives a primary response, including the induction of *Irf1*, *Irf7* and *Irf9* that in turn induces secondary response genes involved in numerous immunomodulatory functions (40, 41). The pathway is also under negative regulation by proteins such as the phosphatase PTPN2 and the negative transcriptional regulators SOCS1 and SOCS3 (42, 43).

Accordingly, we observed that senescence drives an increase in IFNGR1 and IFNGR2 levels, with the former being one of the most prominent changes detected in our proteomics analysis (**Fig. 4A-4C**; Supplementary Fig. S5B). In addition, senescence triggered an increase in several interferon response transcription factors (*Irf1*, *Irf7* and *Irf9* (40, 41)), while causing a concomitant decrease in negative regulators of the pathway’s transcriptional output (*Ptpn2*, *Socs1* and *Socs3* (42, 43)) (**Fig. 4C**). Similar changes were noted in NSP tumor cells treated with different senescence inducers (**Fig. 4C**) and, more broadly, in a panel of 13 human breast, lung, liver and colon derived cancer cell lines triggered to senesce (SENESCOPEDIA) [**Fig. 4D**; Supplementary Fig. S5D-S5G; ref: (37)]. Therefore, changes in components of the Type II IFN signaling apparatus is a broad feature of senescent cells, independent of cell type, cell genotype, species, and nature of the senescence inducer.

**Figure 4.**
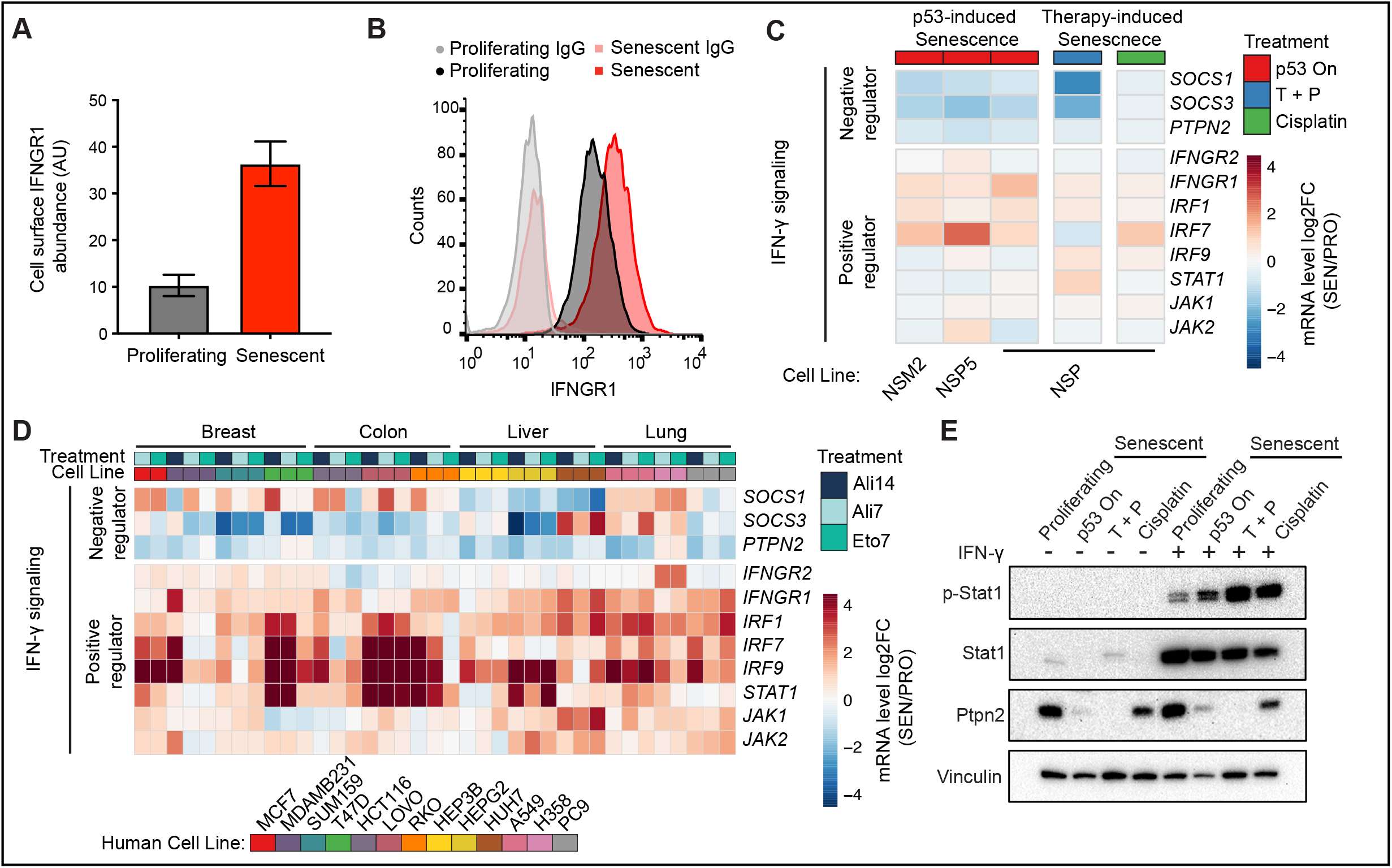
Senescent cells are primed to sense and amplify IFN-γ signaling. A and B, IFNGR1 level on proliferating and senescent cells profiled by mass spectrometry and validated by flow cytometry (B). AU, arbitrary unit. Data is presented as mean ± s.e.m. n = 6 for both proliferating and senescent group. C, Transcriptomic analysis of selected genes regulating IFN-γ signaling from RNA-seq data of 3 independent p53-restorable cell lines (NSP, NSM2, NSP5) restoring p53 along with NSP cells treated with two other senescence triggers. T+P, trametinib plus palbociclib. D, mRNA expression of selected genes involved in IFN-γ signaling in human cell lines triggered to senesce. Treatment: Ali, alisertib; Eto, etoposide; number indicates the length of treatment (days). Data is obtained from the public dataset SENESCOPEDIA (37). E, Immunoblot analysis of NSP cells under different senescent triggers, in presence or absence of IFN-γ (1 ng/ml). p-Stat1, phospho-Stat1 (Tyr701)

The concurrent increase in IFN-γ effectors and decrease in negative regulators led us to hypothesize that senescent cells become primed to sense IFN-γ within their environment. To test this hypothesis directly, we treated proliferating and senescent NSP cells with recombinant IFN-γ at the dose that does not impact their viability (Supplementary Fig. S5H) and performed immunoblotting phosphorylated STAT1. While IFN-γ dramatically increased the base line levels of STAT1 in both states, senescent cells accumulated higher levels of phosphorylated STAT1, irrespective of the senescence trigger (**Fig. 4E**). As predicted from transcriptional analyses, senescence also triggered a decrease in protein levels of the negative regulator of IFN-γ-induced responses, PTPN2 (44), irrespective of the presence of exogenous IFN-γ (**Fig. 4E**). While NSP cells themselves do not produce IFN-γ in either their basal or senescent state (Supplementary Fig. S5I), it was readily detected in tumor tissue protein extracts (Supplementary Fig. S5J), raising the possibility that such amplified IFN-γ signaling in senescent cells may be operative in vivo.

### Senescence and extracellular IFN-γ cooperatively upregulate the antigen processing and presentation machinery

To better understand the functional contribution of IFN-γ sensing to the senescence program, we next performed RNA-seq on proliferating and p53-restored senescent NSP tumor cells treated or not with recombinant IFN-γ (50 pg/mL). While exogenous IFN-γ had no effect on SASP gene expression in either proliferating or senescent conditions (**Fig. 5A**), its impact on certain IFN-γ stimulated genes of senescent cells was profound. Specifically, supervised clustering of the Hallmark “IFN-γ response signature” across proliferating and senescent cells revealed three DEG modules: (i) genes that were downregulated during senescence irrespective of IFN-γ (including the aforementioned negative regulators); (ii) genes that were upregulated during senescence irrespective of IFN-γ and, interestingly, (iii) a substantial set of DEGs that are cooperatively induced by the combination of senescence and IFN-γ (**Fig. 5B**). Therefore, senescence triggers quantitative and qualitative changes in the transcriptional response to IFN-γ.

**Figure 5.**
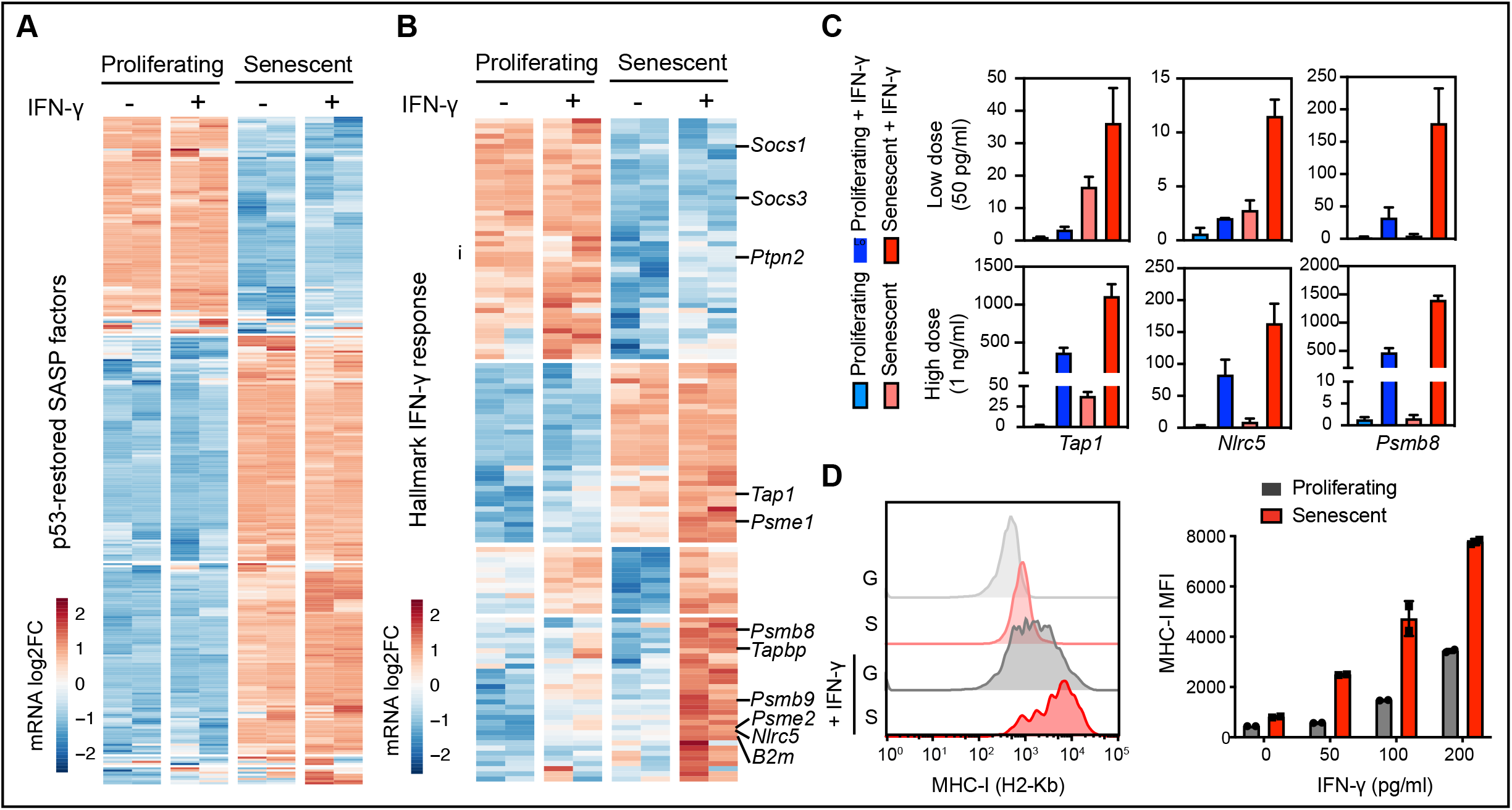
Senescence and extracellular IFN-γ cooperate to upregulate antigen processing and presentation machinery. A and B, mRNA expression of genes in proliferating and senescent NSP cells *in vitro* in the presence or absence of IFN-γ (50 pg/ml) treatment. mRNA level is normalized to the mean expression of the gene in all samples. A, differential expressed genes encoding SASP factors in our model. B, IFN-γ response genes from Hallmark signature database. C, RT-qPCR of selected antigen presentation pathway genes in proliferating and senescent cells treated with low (50 pg/ml) or high (1 ng/ml) concentration of IFN-γ. Samples are from 2 biological replicates. D, MHC-I level of proliferating and senescent cells treated with IFN-γ for 24 hours measured by flow cytometer. MFI, median fluorescence intensity. Data is presented as mean ± s.e.m.

One well-established output of IFN-γ signaling regulating cells’ susceptibility to adaptive immune surveillance is an increased capacity for antigen presentation mediated by MHC class I molecules (MHC-I) (40, 45) and, indeed, many of the genes upregulated in senescent cells (class ii genes) or hyper-induced in the presence of exogenous IFN-γ (class iii genes) included components of the antigen presentation machinery. Among the genes induced during senescence (class ii genes) were *Tap1*, transporters associated with antigen processing, and *Psme1*, a proteosome factor associated with antigen processing (46). Those hypersensitive to exogenous IFN-γ (class iii genes) included *Nlrc5*, a transcriptional co-activator of MHC-I genes (47), the MHC-I assembly factor *Tapbp*, and the MHC-I subunit *B2m*. Two other genes in such class were components of the immunoproteasome (*Psmb8, Psmb9*), whose actions can alter the repertoire of presented peptides when overexpressed and are associated with an improved tumor response to immune checkpoint blockade (48). These synergistic effects of senescence and IFN-γ on the expression level for several genes were confirmed by RT-qPCR and were retained at even higher levels of exogenous IFN-γ (**Fig. 5C**).

In line with the above observations, senescent cells responded to low levels of exogenous IFN-γ by more robustly upregulating MHC-I. Hence, while cell surface levels of MHC-I were low in both proliferating and senescent cells at baseline (Supplementary Fig. S5K and S5L) and induced by exogenous IFN-γ, senescent cells showed a significant increase of the mean fluorescence index in MHC-I expression compared to proliferating controls (**Fig. 5D**). Similar synergies were observed for cell surface HLA expression (identical to MHC-I in mice) in human liver cancer cells triggered to senescence with nutlin, which engages a p53-dependent senescence program (49), or trametinib/palbociclib, which preferentially targets tumor cells with an activated MAPK pathway (20) (Supplementary Fig. S6A-S6C). Of note, the combinatorial effects of drug treatment and IFN-γ on HLA expression required senescence induction and were lost in liver tumor cells that failed to senesce, either due to the cells harboring spontaneous or engineered p53 mutations (irresponsive to nutlin) or lacking an activated MAPK pathway (irresponsive to trametinib/palbociclib). These data imply that murine and human cells triggered to senesce acquire an increased capacity for antigen processing and presentation in the presence of limiting quantities of IFN-γ.

### Senescent tumor cells hyperactivate the IFN-γ signaling pathway in vivo

To determine the *in vivo* consequences of the rewiring of IFN-γ signaling identified in senescent cells, we next adapted an IFN-γ sensing (IGS) reporter system to directly visualize in intracellular IFN-γ signaling activation in real time (50). This reporter consists of a series of interferon gamma-activated sequences, followed by a cDNA sequence encoding ZsGreen1 fluorescent protein and is linked to a constitutively expressed RFP transgene to visualize transduced cells (**Fig. 6A**). NSP tumor cells expressing this construct were RFP positive and showed a dose-dependent increase in ZsGreen1 signal upon treatment with IFN-γ in vitro that increased following p53 induction or following treatment with senescence-inducing drugs (**Fig. 6B**; Fig. S7A and S7B).

**Figure 6.**
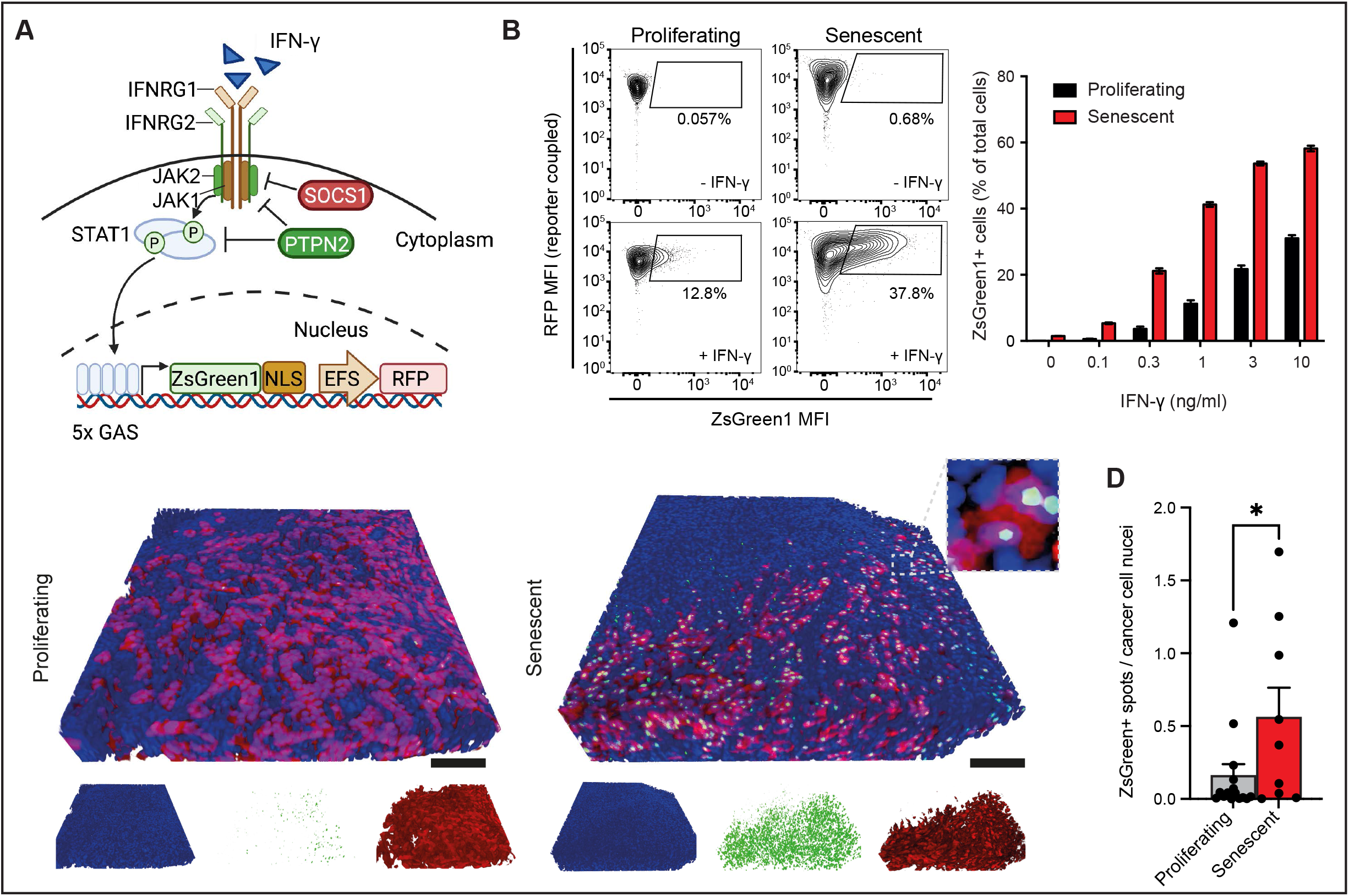
Senescent tumor cells show a hyperactivation of IFN-γ signaling. A, Graphic illustration of IFN-γ sensing (IGS) reporter. B, Left panel, representative flow cytometry plots measuring ZsGreen1 signals in proliferating and senescent NSP cells treated with 1 ng/ml IFN-γ. Right panel, quantification of the percentage of ZsGreen1 positive cells upon IFN-γ treatment. C and D, Representative 3D imaging of tissue cleared tumors from the orthotopically injected liver NSP cell line expressing IGS reporter (C). Quantification of 3 randomly selected fields from the liver tumor of each mouse (D). N=5 and N=3 for the proliferating and senescent group respectively. MFI, median fluorescence intensity. Data is presented as mean ± s.e.m. Two-tailed student t-test is used. *p < 0.05. Scale bar, 100 µm.

Having validated that the IGS reporter responds to IFN-γ *in vitro*, we next used this system to monitor signaling activity following senescence induction in tumors. RFP-positive tumor cells (on Dox) were injected into the livers of dox-fed syngeneic recipients and, upon tumor manifestation, doxycycline was removed to induce p53 expression and trigger senescence as above (see Figures 1 and 2). Tumors were isolated from these animals during the phase of tumor regression (10 days post Dox withdrawal) and subjected to the 3D imaging after tissue clearing protocol to assess the pattern of the ZsGreen1 reporter in comparison to proliferating tumors (maintained on Dox). While proliferating tumor cells showed little, if any, reporter expression, tumor cells in the senescent state displayed a prominent ZsGreen1 signal (**Fig. 6C and 6D**; Supplementary video S2). Although the altered composition of immune cells in tumors following senescence induction may contribute to this enhanced signal, in vitro assays allowing for normalization of tumor cell:activated T cell ratios still showed a significant increase in the ZsGreen1 reporter signal in senescent tumor cells as compared to proliferating controls (Supplementary Fig. S7C). These data imply that the enhanced IFN-γ signaling occurring within tumors upon senescence induction is not merely a consequence of more immune cell recruitment but also of a rewired IFN-γ sensing tumor program that is associated with enhanced tumor immune surveillance in vivo.

### IFN-γ signaling in senescent tumor cells is necessary for immune surveillance

Our results imply that the immune-mediated clearance of senescent NSP tumor cells involves the combined effects of SASP, known to stimulate immune cell recruitment (10,20,21), together with a previously underapreciated enhanced ability of senescent cells to sense and respond to extracellular signals, as an example shown here with IFN-γ. To test the requirement of the senescence-activated IFN-γ sensing program for tumor cell clearance, we examined the effects of cell-intrinsic IFNGR disruption, or IFN-γ depletion in the host, on the clearance of NSP tumor cells triggered to senesce in vivo. Indeed, tumor regressions following p53 induction and senescence were blunted upon knock-out of IFNGR1 (**Fig. 7A and 7B**; Supplementary Fig. S8A-S8B), a defect that was even more pronounced for IFNGR-intact tumors engrafted into *Ifng-/-* mice (**Fig. 7C and 7D**; Supplementary Fig. S8C). Importantly, this impaired senescence surveillance phenotype was not simply a result of lack of immune cell recruitment, as CD45 positive immune cells (including F4/80 macrophages and CD8 T cells) were efficiently recruited into tumors harboring senescent *Ifngr1-/-* tumor cells that were unable upregulated MHC-I (**Fig. 7E and 7F**; Supplementary Fig. S8D-S98F). In agreement, co-culture assays controlling for the degree of exposure of NSP cells to the immune cell types mediating senescence surveillance in vivo (Fig. 2E,F) recapitulated the expected IFNGR-dependent increase in immune-mediated lysis of senescent tumor cells –a dependence that was not observed in proliferating counterparts under the same conditions (Supplementary Fig. S9). Taken together, these data indicate that the enhanced sensing capacity of senescent tumor cells for IFN-γ in the environment acts in concert with SASP-simulated immune cell recruitment and enable tumor cell surveillance, leading to potent tumor regressions.

**Figure 7.**
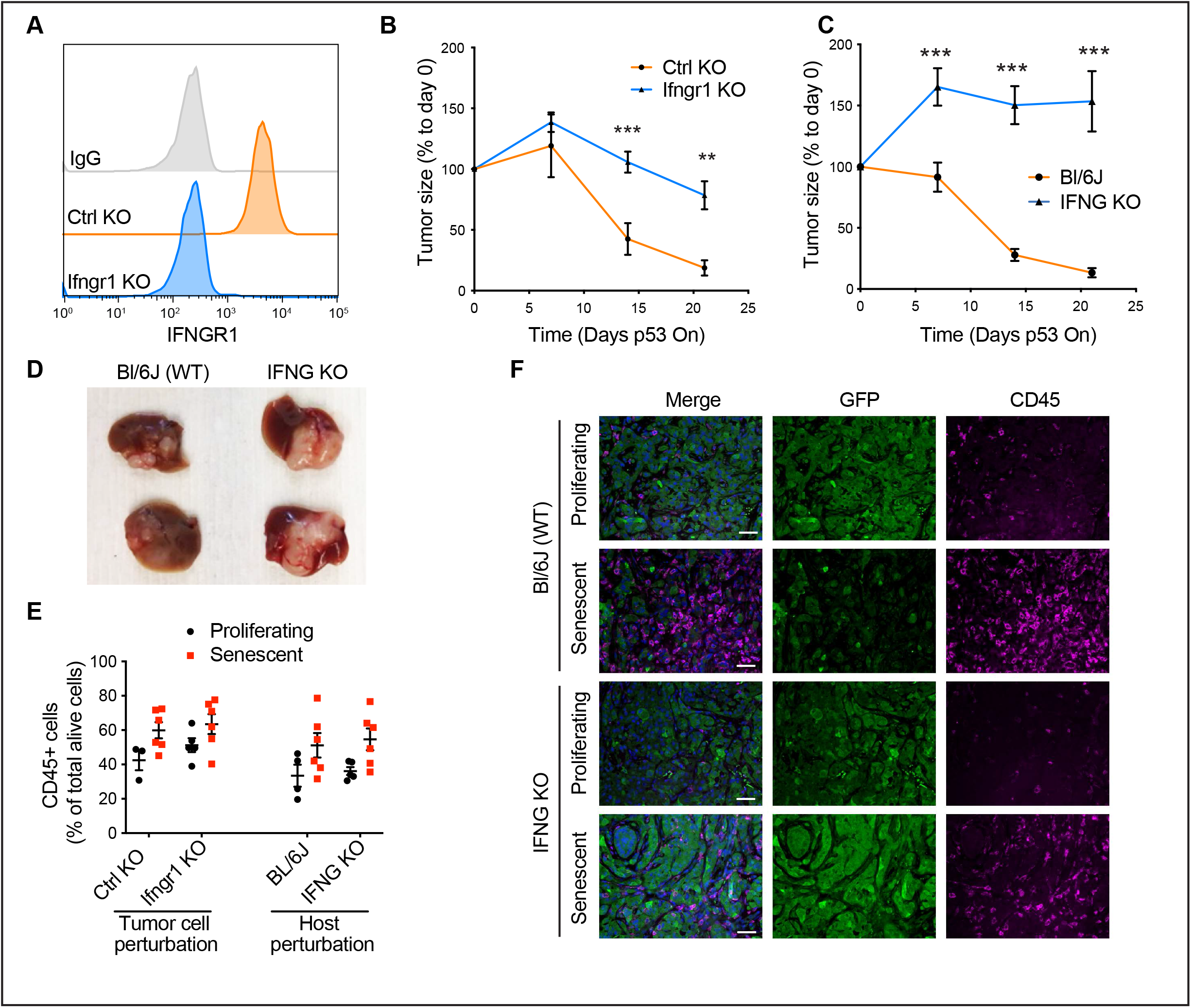
IFN-γ signaling in senescent tumor cells is necessary for immune surveillance. A, *Ifngr1* knockout (KO) validated by flow cytometry. A control sgRNA targeting a gene desert located on Chr8 (Ctrl KO) serves as a control. B, Tumor regression phenotype of Ifngr1 KO or control sgRNA-transfected tumor cells orthotopically injected into Bl/6N mice upon p53 restoration. C, Tumor regression phenotype of parental NSP tumor cells orthotopically injected into WT or *Infg* KO mice upon p53 restoration. D, Representative macroscopic images of tumor collected at day 21 after p53 restoration from (C). E, Flow cytometry analysis of CD45 abundance in tumor from indicated groups. F, Representative immunofluorescence in p53-suppressed (proliferating) and p53-restored (senescent, 7 days after p53 restoration) tumor from the indicated host. NSP tumor cells were transduced with GFP-expressing vector for visualization. Scale bar, 50 µm Data is presented as mean ± s.e.m. Two-tailed student t-test is used. *p < 0.05; **p < 0.01; ***p < 0.001.

## DISCUSSION

Enabled by a murine tumor model in which tumor cell proliferation and immune evasion versus tumor cell senescence and CD8 T cell-mediated tumor regression are under tight genetic control, we reveal how senescent cells dramatically alter their ability to both send and receive environmental signals (Supplementary Fig. S10). Consistent with other senescence programs, p53-driven senescence induction led to the silencing of proliferative genes and induced SASP. However, we also observed a profound effect on the expression of plasma membrane encoding genes including a range of growth factor receptors and cytokine receptors that are predicted to drastically alter how senescent cells respond to environmental signals. Importantly, a similar shift in the expression of cell surface sensors was observed in a broad range of murine and human tumor cells treated with senescence inducing agents, implying that the altered sensing program is a general hallmark of the senescent state.

One of the prominent sensing pathways altered in senescent cells involves type II IFN signaling. In our model and across all senescent states we examined, senescence is accompanied by transcriptional and protein expression changes predicted to enhance pathway signaling output in the presence of exogenous IFN-γ. Indeed, senescent cells more robustly activated IFN-γ effectors in response to IFN-γ in vitro and in vivo, and the presence efficient CD8 T cell-mediated clearance of senescent tumor cells required an intact IFN-γ effector pathway and the presence of IFN-γ in the environment. While pathway analysis of the senescent state transcriptome invariably identified type I and type II IFN signaling as enriched features, the fact that many components of these pathways overlap and that IFN-γ is rarely observed in SASP has left mechanistic questions regarding type II IFN signaling in senescence biology largely unexplored. Our studies demonstrate that the presence of “IFN Gamma Response” in pathway analysis of senescent signatures obtained in cell culture reflects their altered capacity of IFN-γ sensing whose output is most prominent in vivo.

Perhaps the most well-established output of type II IFN signaling involves its ability to induce the antigen presentation machinery and, indeed, IFN-γ induced cell surface expression of MHC-I (or HLA in human cells) in our model under both proliferating and senescence conditions. However, MHC-I upregulation was more pronounced in senescent cells, an effect that correlated with increased expression of the transporter associated with antigen processing and the synergistic effects of exogenous IFN-γ on yet other antigen processing factors and structural components of MHC-I. A similar hypersensitivity to IFN-γ in inducing MHC-I/HLA levels was observed in human cancer cell lines triggered to senesce in vitro as well as in murine cells following senescence induction in vivo. These results imply that the senescence program can enhance the ability of non-immune cell types to present antigen, and thereby facilitate immunosurveillance.

Our results support a model whereby the ultimate impact of senescent cells on tissue biology is dictated by the combined effects of how they send and receive environmental signals. On one hand, senescent cells induce the SASP, which triggers tissue remodeling and alters the cell state and composition of immune cells in the environment. On the other, senescent cells dramatically alter their surfaceome leading to a differential ability to sense environmental factors, herein exemplified by IFN-γ. Importantly, disruption of IFN-γ signaling in the tumor cells had no effect on senescence induction or immune cell recruitment yet impaired tumor regression, indicating that altered environmental sensing acts in concert with SASP to determine the ultimate output of the senescence program – in this case, immune surveillance. These effects appear to be part of a coordinated epigenetic process, as both the SASP and sensing programs show a prominent dependence on the chromatin remodeling factor Brd4.

While immune surveillance in our model involves CD8 T cell-mediated targeting, other innate or adaptive immune cell types can recognize and clear senescent cells in different contexts or, alternatively, surveillance may not occur at all (18, 19). Undoubtedly, some of these distinctions reflect the secretion of different SASP factors (13, 14), though our results raise the possibility that the extent and nature of altered environmental sensing may also contribute to heterogeneity in senescence biology. Certainly, the fact that senescent cells can respond differently to environmental signals implies that their ultimate molecular state in tissues will be different than in cell culture, highlighting the need to better characterize the process in vivo.

Our results may help understand the paradoxical effects of senescence biology in physiology and disease and have implications for the effective use of senescence-modulating therapeutics. For example, in our model, the difference between senescence cell clearance and persistence was determined by the presence of environmental IFN-γ and the integrity of the type II IFN signaling in the tumor cells. This suggests that variation in the ability of senescent to recruit and sense IFN-γ secreting immune cells or other immune cell types could profoundly affect the tumor suppressive and regenerative properties associated with senescent cell clearance that, if impaired, could enable senescent cell persistence and tissue decline. In the context of cancer, therapies that induce tumor cell senescence – a cytostatic program – sometimes trigger immune-mediated tumor regression. As such, heterogeneity in the SASP (which can vary between tumor cell types and senescence inducers) or IFN-γ sensing and output (perhaps affected by deletion or mutation of IFN-γ pathway or HLA components (51) or the reversible transcriptional mechanisms uncovered here) may influence the effectiveness of such therapies in patients. By contrast, strategies to enhance the immune surveillance of senescent cells by increasing their sensitivity to IFN-γ (e.g. with PTPN2 inhibitors), or through immune checkpoint blockade or CAR T therapy, may help bias program output towards tumor cell rejection.

## SUPPLEMENTARY FIGURE LEGENDS

**Supplementary figure 1.**
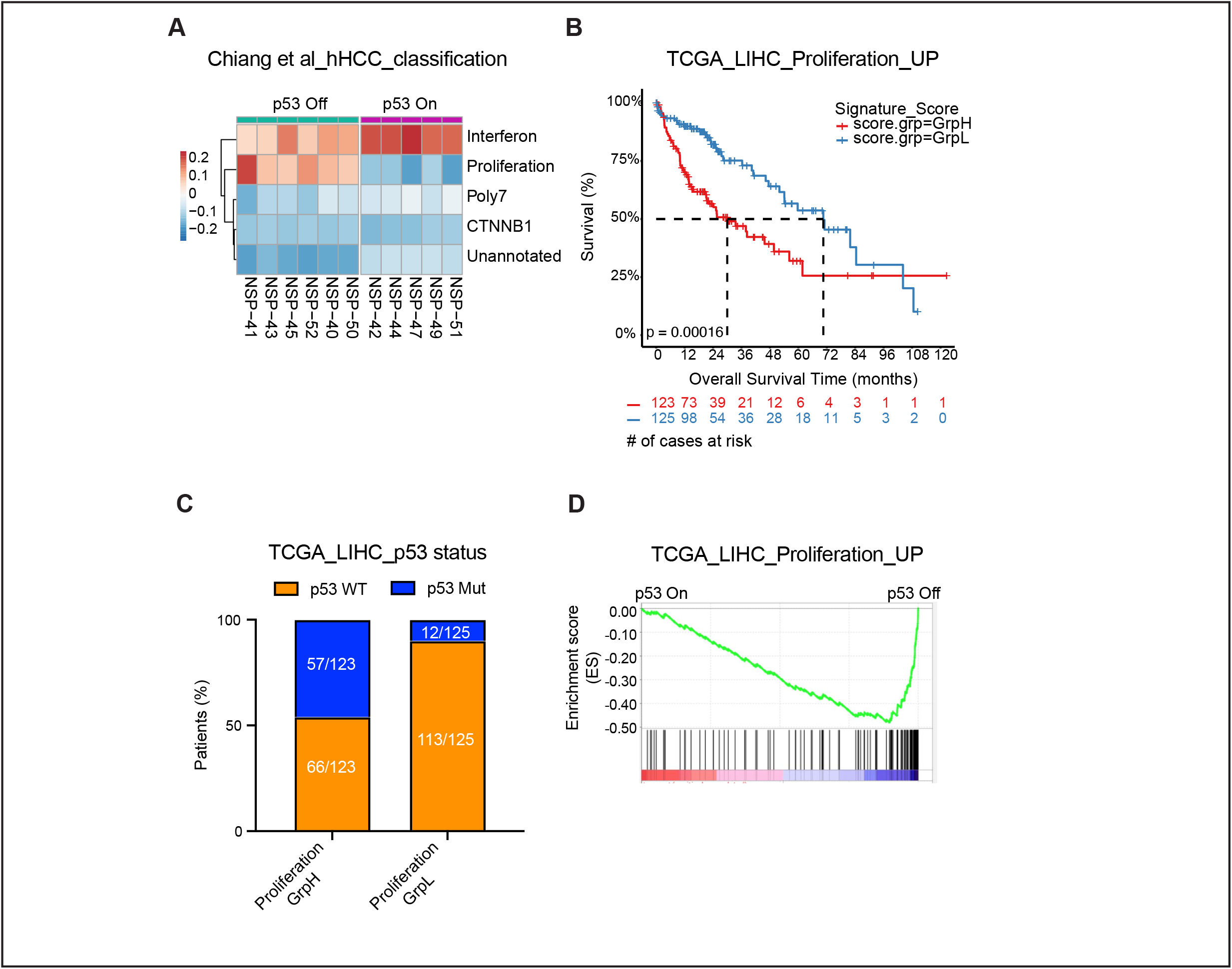
The p53-restorable immunocompetent liver cancer model resembles “Proliferation class” in human hepatocellular carcinoma. A, Classification of Nras-driven, p53-restorable HTVI liver tumors using bulk RNA-seq using signatures from publicly available dataset. B, TCGA data of liver cancer patients stratified by Proliferation Class signature score and grouped into top (GrpH) 33% and bottom (GrpL) 33%. C, p53 mutation status of liver cancer patients from TCGA, grouped by Proliferation Class signature score in (B). D, Gene Set Enrichment Analysis between p53 On and p53 Off HTVI liver tumor using gene signature upregulated in Proliferation Class.

**Supplementary figure 2.**
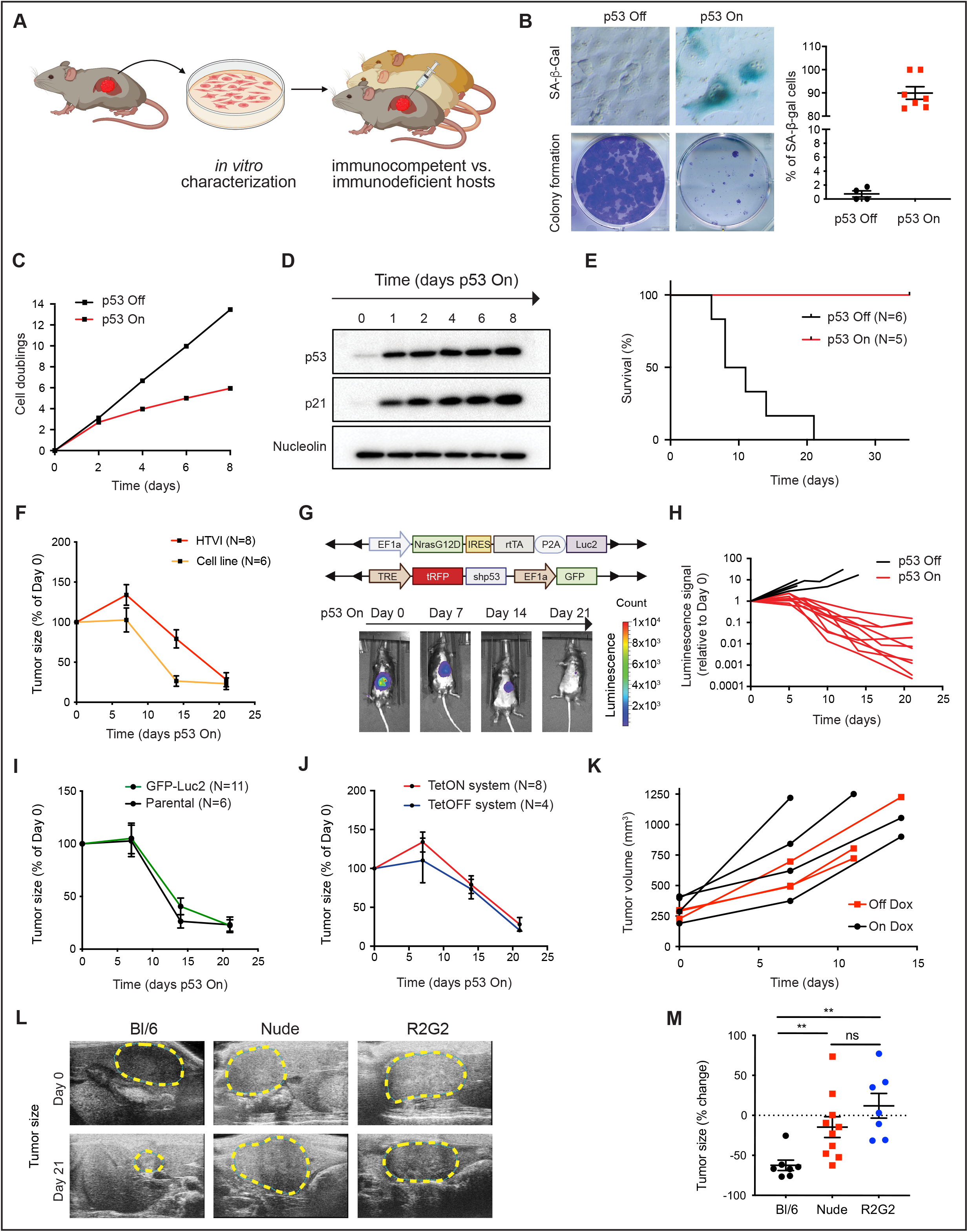
Establishment of a genetically controlled tumor-specific senescence mouse model. A, Graphic illustration of cell lines generation from the HTVI model and subsequent in vivo tumor cells orthotopic injection experiment. B, *In vitro* characterization of p53-restorable liver cancer cell line, NSP, by colony formation assay and SA-β-Gal staining. Several line has been generated and NSP line is predominantly used for the *in vitro* and orthotopic injection study. C, Comparison of cell doublings in p53 On and p53 Off tumor cells. Experiment is performed in triplicate wells. D, Immunoblot analysis of p53-restorable tumor cells cultured *in vitro*. E, Survival analysis of mice from orthotopic injection model of NSP tumor cells. F, Comparison of tumor regression phenotype between HTVI and orthotopic injection model measured by ultrasound. G, Representative bioluminescence images of tumor regression upon p53 restoration in the HTVI model using luciferase-containing transposon constructs. H, Longitudinal tracking of tumor growth in orthotopic injection model. NSP cells were transduced with a GFP-luciferase vector to enable bioluminescence imaging. Each line represents an individual mouse. Related to Supplementary. Fig S1G. I, Comparison of tumor regression phenotype upon p53 restoration between parental and GFP-luciferase transduced tumor cells in orthotopic injection model. J, Comparison of tumor regression phenotype upon p53 restoration between Tet-On (rtTA) and Tet-Off (tTA) system in the HTVI model. K, Comparison the growth of tumor harboring a p53 hairpin driven by a constitutive promoter in the presence or absence of doxycycline (Dox) treatment. Each line represents an individual mouse. L, Representative ultrasonogram of the tumor size at indicated time after restoring p53 expression in immunocompetent and -deficient mice. M, Quantification of tumor size change between day 7 and 14 after p53 restoration from mice shown in (L) showing a trend to a greater defect of tumor regression in R2G2 compared to nudes. Data is presented as mean ± s.e.m.

**Supplementary figure 3.**
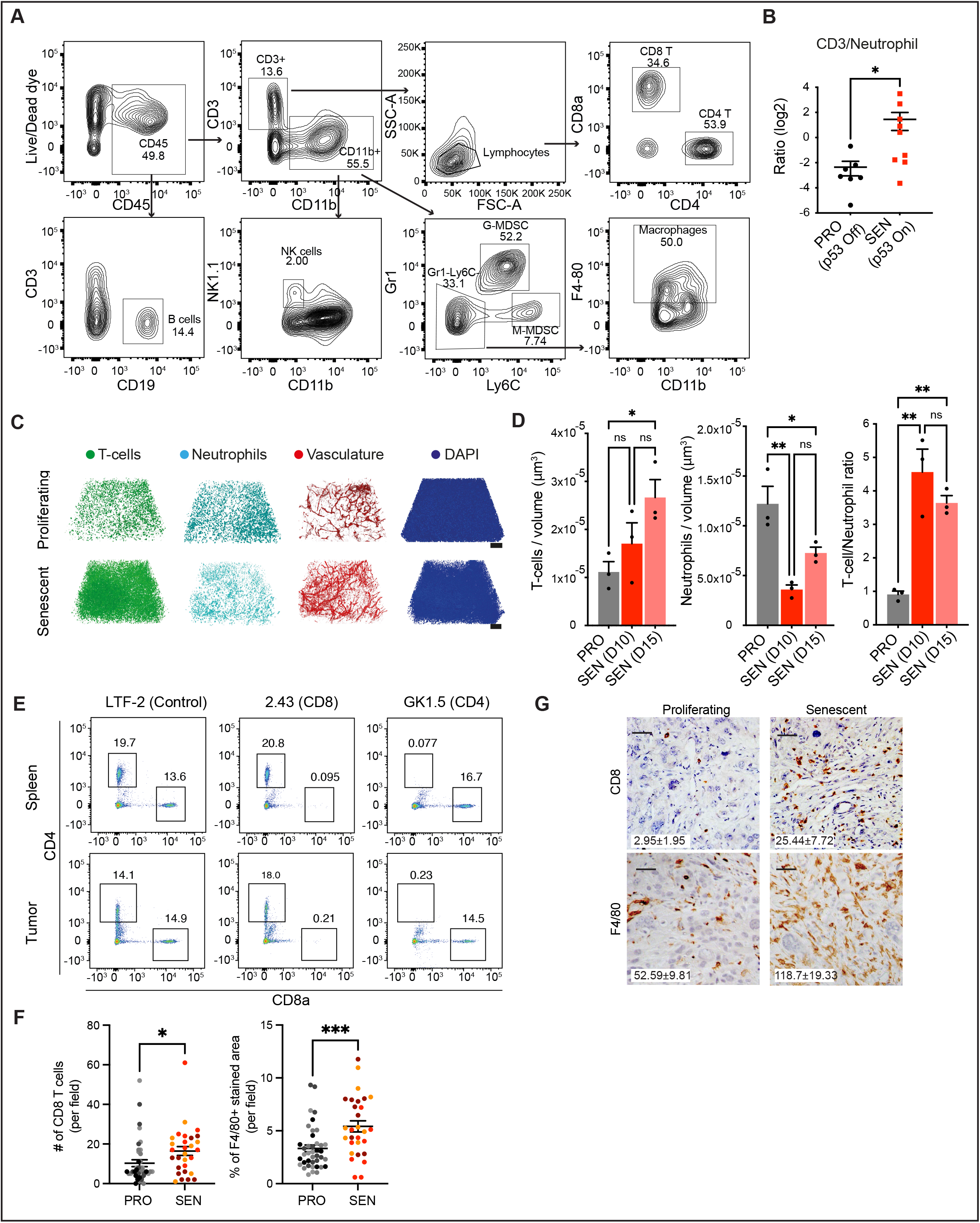
Senescence engagement switches from immune-suppressive to immune-activated tumor microenvironment. A, Gating strategy for immunophenotyping. Related to Fig. 2B. B, CD3 to neutrophil ratio calculated from flow cytometry measurements in Fig. 2B. C, Individual channels of 3D imaging after tissue clearing from Fig. 2D D, Quantification of CD3 T cells and neutrophil density and the CD3/neutrophil ratio at indicated time points of tumor collection (D, day). PRO, proliferating. SEN, senescent. Related to Fig. 2D. E, representative flow cytometry plots of CD4 and CD8 T cells depletion through corresponding antibodies. F, Quantification of the number of CD8 T cells and the percentage of F4/80+ area per field in NSP tumor. 9-10 random fields are taken per mouse sample and each color represent a mouse. N=4 and 3 of the mice harboring p53-suppressed (PRO) or -restored tumor (SEN) respectively. Related to Fig. 2F. G, Representative IHC images of CD8 T cells and F4/80 positive macrophages staining in the HTVI-generated tumor. p53-restored tumors (senescent) were collected 14 days after randomization to indicated treatmentf. Quantification of number of cells per field from 2-3 random fields each mouse and N ≥ 4 mice per group. Scale bar, 100 µm. Data is presented as mean ± s.e.m. A two-tailed student t-test is used. **p < 0.01, *p < 0.05.

**Supplementary figure 4.**
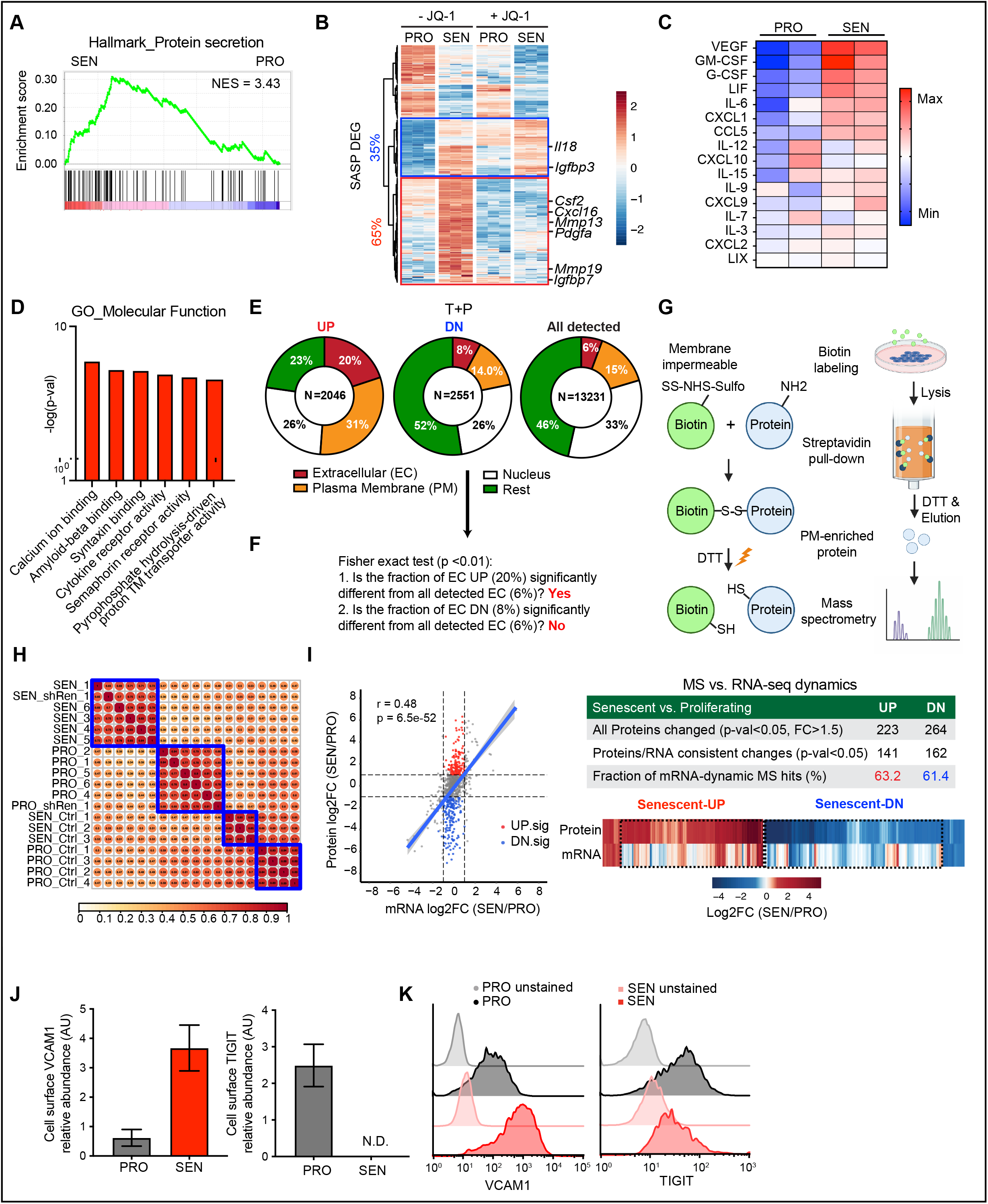
Cell surfaceome is substantially remodeled in senescent cells. A, GSEA (Hallmark) of RNA-Seq data from proliferating (PRO, p53 Off) vs. senescent (SEN, p53 On for 8 days) NSP liver tumor cells in vitro. B, Transcriptomic analysis of differential expressed genes (DEGs) encoding secretory factors (SASP factors) in NPS cells in the presence or absence of JQ-1 treatment. C, Cytokine array of conditioned medium collected from proliferating and senescent (p53 On Day 6 to Day 8) cells in vitro. Samples are from 2 independent biological replicates. D, GO analysis of DEGs encoding plasma membrane proteins upregulated in senescent cells. Related to Fig. 3C E, Subcellular localization of detected DEGs (TPM > 1; p < 0.05; fold change > 2) from in vitro NSP cells treated with trametinib/palbociclib (T+P) or vehicle. F, Calculation of p value of Fisher exact test using data from the supplementary Fig. S4E. A more stringent criteria for statistic significance is used, p < 0.01. Related to Fig. 3E. G, Graphic illustration of the protocol of plasma membrane-enriched mass spectrometry (MS). H, Correlation plot of MS samples. I, Left panel, XY plot of total proteins profiled by MS against corresponding transcriptomic expression profiled by RNA-seq. Right panel, summary of MS profiling and RNA-seq comparison J and K, validation of two MS hits using flow cytometry. PRO, proliferating. SEN, senescent. N.D., not detected. Data is presented as mean ± s.e.m.

**Supplementary figure 5.**
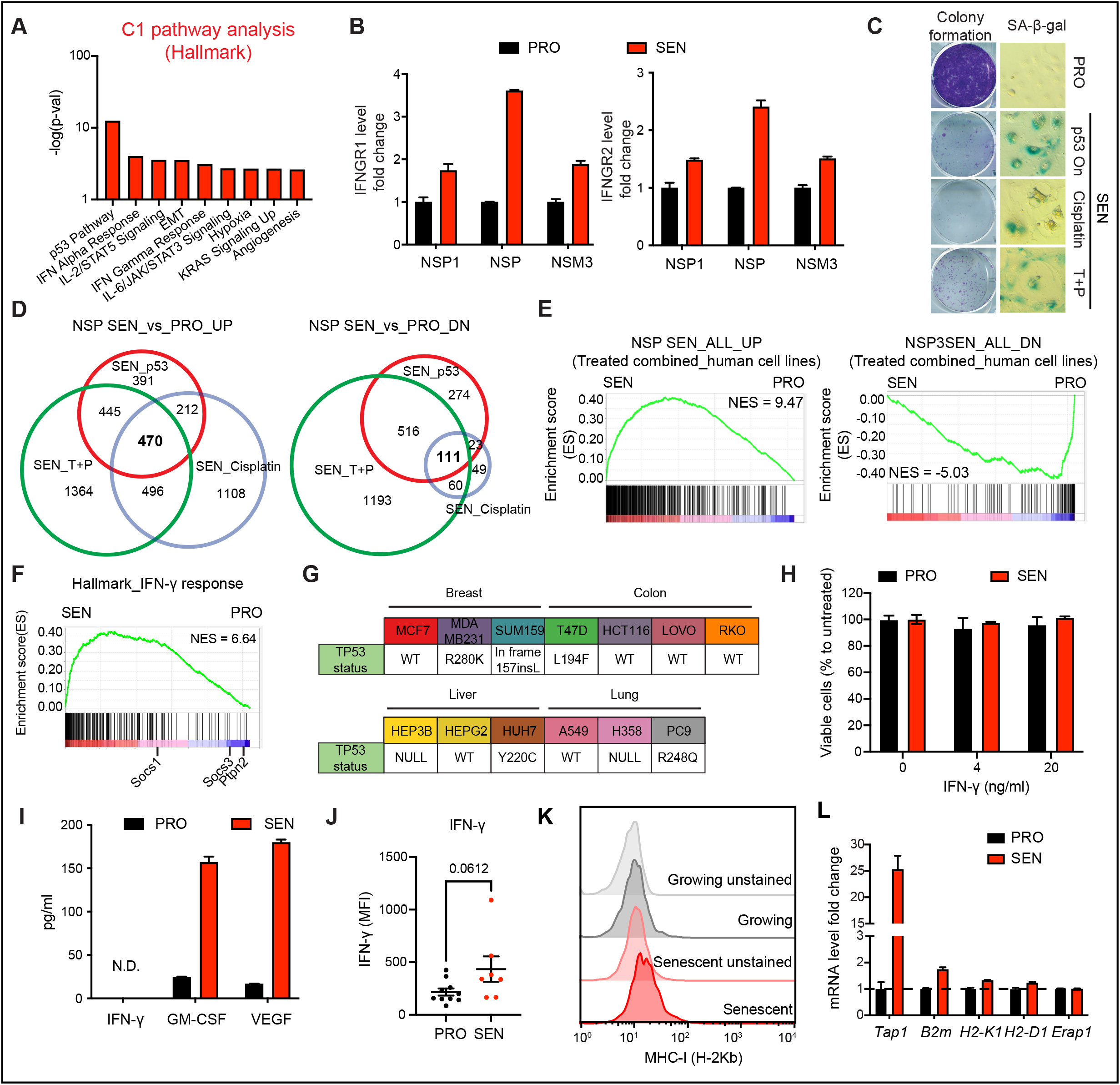
Sensitization to IFN-γ in senescent cells is independent of p53 status. A, Pathway analysis of cluster 1 (C1, senescence-specific) shown in Fig. 3D against MsigDB Hallmark genesets. B, IFNGR1 and IFNGR2 level validated in 3 independent p53-restorable liver cancer cell lines. NSP is predominantly used in this study. C, SA-β-gal staining of NSP cells treated with different senescence triggers. D, Overlapping DEGs from RNA-seq in NSP cells treated with different senescence triggers to identify common signatures upregulated (UP) and downregulated (DN) in proliferating (PRO) vs. senescent (SEN) cells, which is composed of 111 and 470 genes respectively. p53, p53 restoration; T+P, trametinib plus palbociclib. E, GSEA of the combined RNA-seq results from all human cell lines trigger to senesce in SENESCopedia showing enrichment of genes upregulated (ALL_UP) or downregulated (ALL_DN) in our common senescence signature from (D). F, GSEA of same RNA-seq results in (E) showing an enrichment in MSigDB Hallmark IFN-γ response pathway in senescent cells. G, p53 mutation status of human cell lines used in SENESCopedia. H, Viability assay of proliferating and p53-restored senescent NSP cells treated with indicated dose of IFN-γ for 48 hours. I, Cytokine array of conditioned medium collected from proliferating and senescent (p53 On Day 6 to Day 8) cells in vitro. Same data collected from Fig. 3C. J, Cytometric bead array (CBA) assay for IFN-γ level from *in vivo* tumor tissue lysate samples. K, MHC-I (H2-Kb) level of proliferating and senescent NSP cells *in vitro*. L, RT-qPCR of selected antigen presentation pathway genes in proliferating and senescent NSP cells *in vitro*. Data is presented as mean ± s.e.m. Two-tailed student t-test is used.

**Supplementary figure 6.**
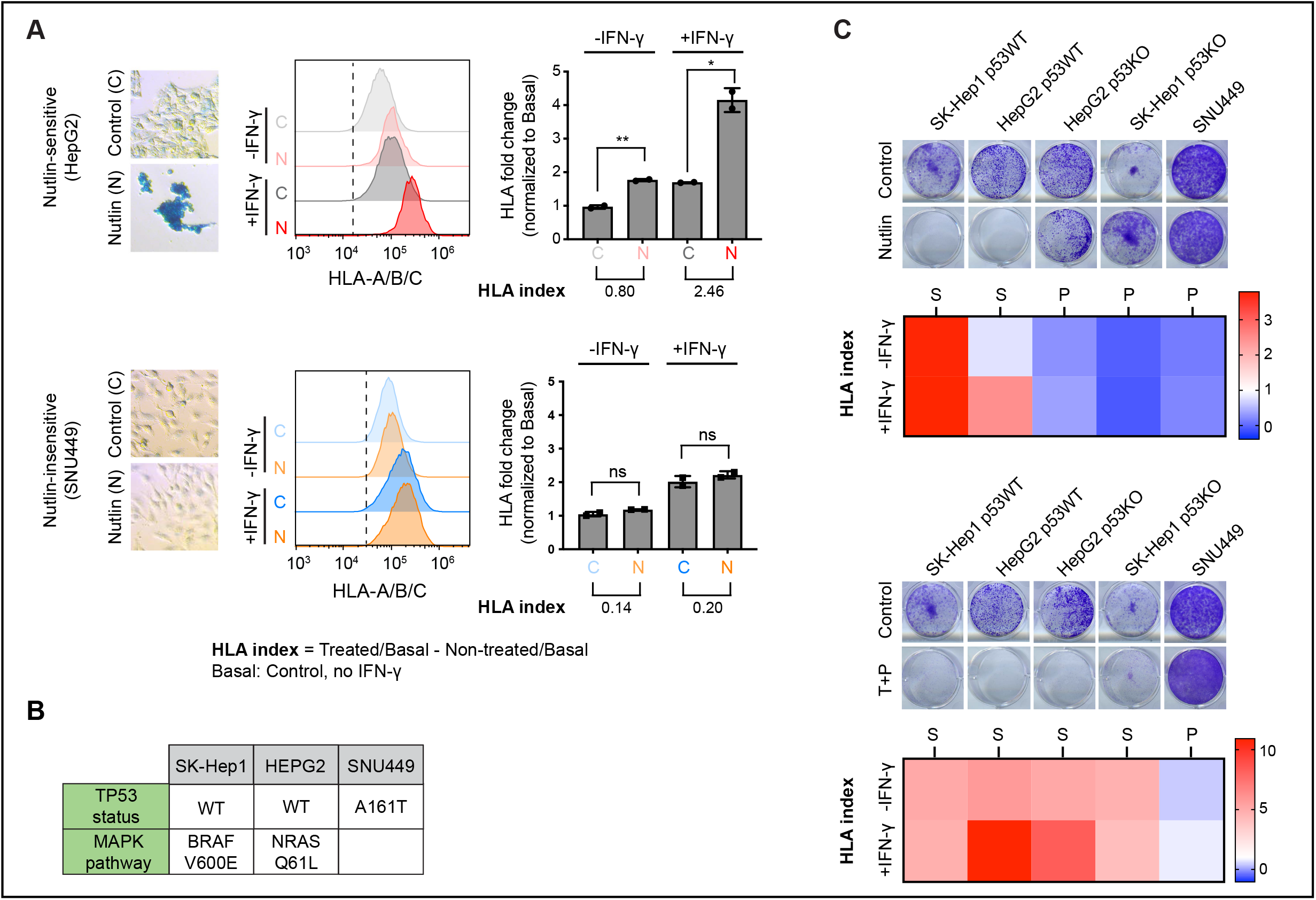
Validation of the cooperativity between senescence and IFN-γ signaling in inducing surface expression of HLA in human cell lines. A, Human liver cancer cell lines and isogenic p53 KO clones were treated with indicated drugs to induce senescence. Cells were treated with human recombinant IFN-γ (1 ng/ml) and HLA-A/B/C was measured after 24h of treatment. “HLA-index” was determined by calculating the HLA level difference between drug-treated vs. untreated cells, in the presence or absence of IFN-γ treatment. Here shown is an example of HLA-index calculation. B, p53 and RAS pathway mutation status of the human cell lines used in this experiment. C, The summary of 5 human cell lines (including two isogenic p53 KO clones) treated with nutlin or trametinib + palbociclib (T+P). S, senescent; P, proliferating. Data is presented as mean ± s.e.m.

**Supplementary figure 7.**
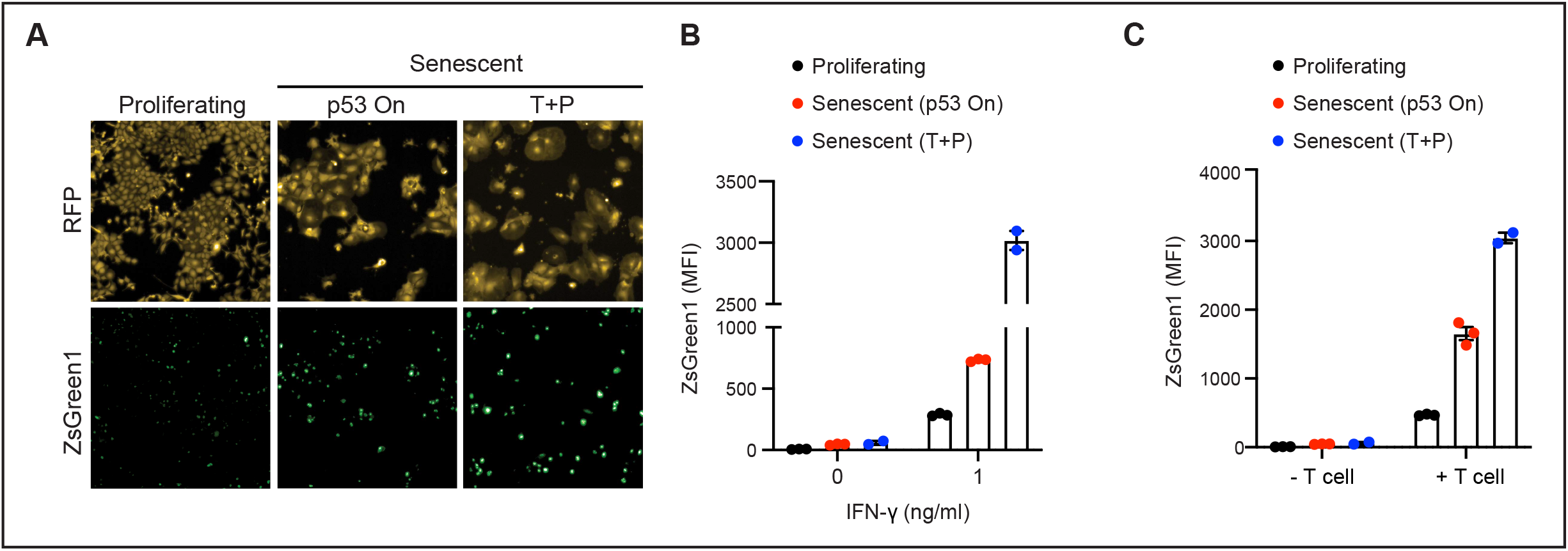
Increased sensitivity of IFN-γ across different senescent triggers in IGS-expressing tumor cells. A, Representative microscopic images of IGS reporter-expressing proliferating and senescent NSP tumor cells triggered by p53 restoration or trametinib + palbociclib (T+P) in combination with IFN-γ (1 ng/ml) treatment. B, Quantification of ZsGreen1 intensity from (A). C, Quantification of ZsGreen1 intensity of NSP tumor cells in the OT-I T cells and SIINFEKL-expressing tumor cells co-culture experiment (E/T ratio 5:1) after 20 h of co-culture. Signal measured by flow cytometry. T+P, trametinib plus palbociclib. MFI, median fluorescence intensity. Data is presented as mean ± s.e.m.

**Supplementary figure 8.**
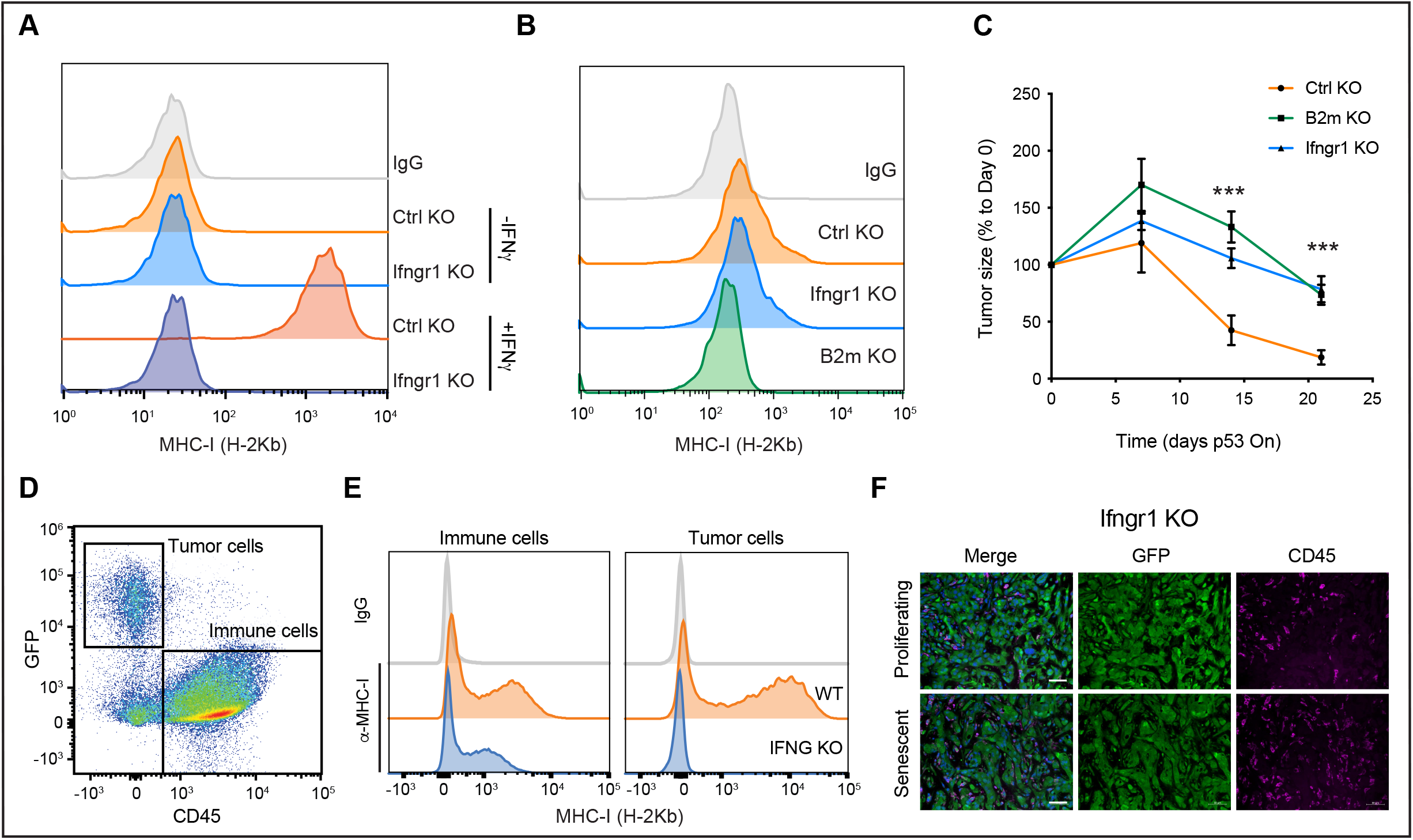
Blunting IFNGR1/IFN-γ signaling in tumor cells phenocopies B2m knockout in senescence surveillance. A, Flow cytometry analysis of MHC-I level in Ifngr1 KO and control sgRNA (Ctrl KO) tumor cells treated with IFN-γ (1ng/ml). B, Flow cytometry analysis of MHC-I level in Ctrl, Ifngr1 and B2m KO tumor cells at the basal level. C, Tumor regression phenotype of Ctrl, Ifngr1 and B2m KO tumor upon p53 restoration. Figure was overlay with Fig. 7B. D, Representative flow cytometry plots gating the GFP+ tumor cells and CD45+ immune cells. Related to Fig. 7E. E, Representative flow cytometry plots showing MHC-I level in tumor cells and immune cells from WT and IFNG KO mice. F, Representative immunofluorescence of Ifngr1 KO tumor in Bl/6N mice. Related to Fig. 7F. Scale bar, 50 µm Data is presented as mean ± s.e.m. Two-tailed student t-test is used. *p < 0.05; **p < 0.01; ***p < 0.001.

**Supplementary figure 9.**
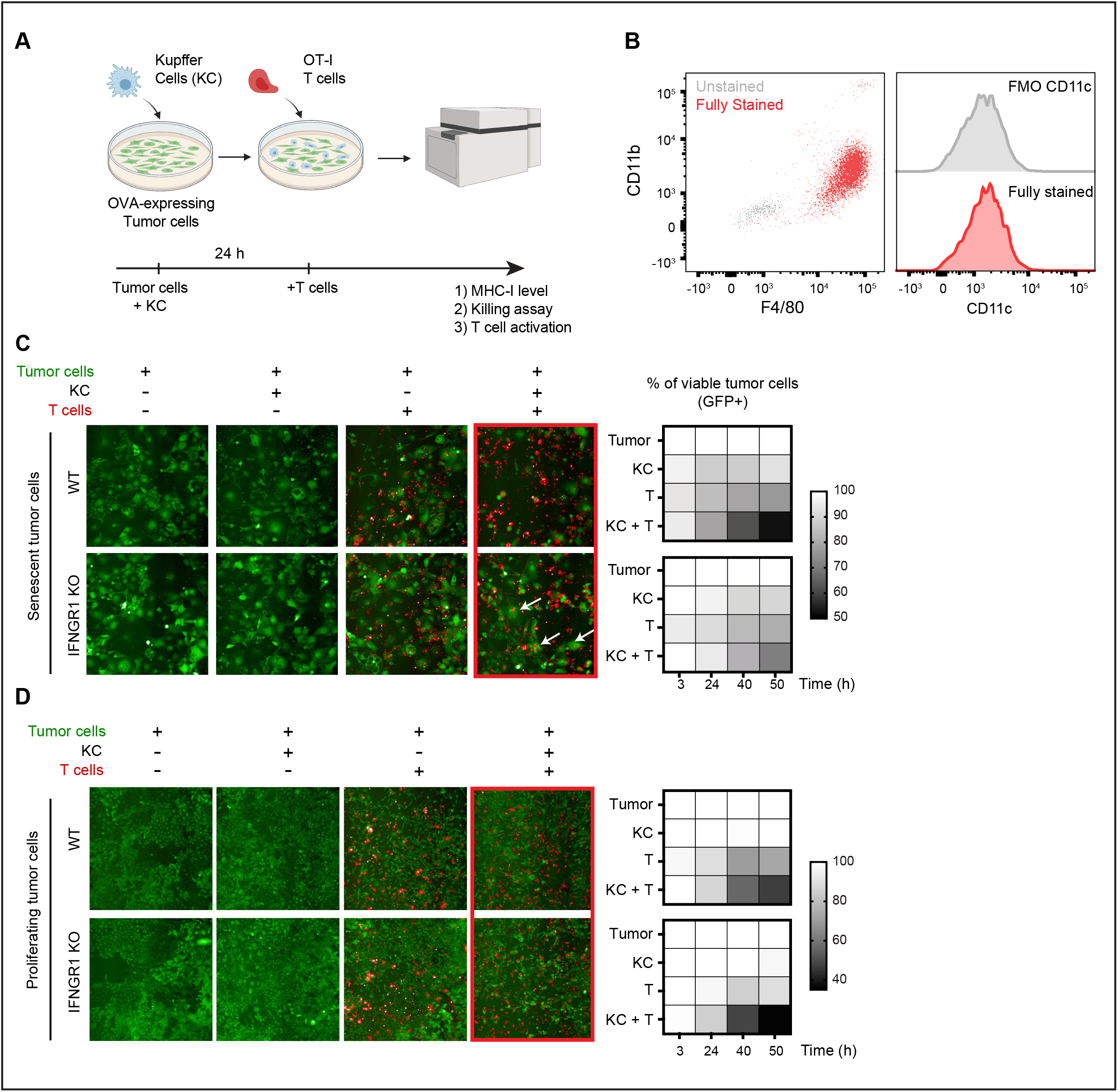
CD8 T cells and macrophages cooperate to kill senescent tumor cells in a IFN-γ -dependent manner. A, Schematic of co-culture experimental setup. OT-I T cells (T), Kupffer cells (KC) and SIINFEKL-expressing NSP cells. B, Purity of isolated KC cells examined by flow cytometry. C, D, Representative time-lapsed images (left) and killing quantification (right) of co-culture assay. Representative images correspond to the end time point (50 hours), and arrows highlight the persistence of senescent (but not proliferating) tumor cells perturbed for IFNGR1 signaling. The quantification of viable senescent (SEN, C) or proliferating (PRO, D) cells is calculated by measuring the changes of GFP positive cell number normalized to the untreated (no immune cells) control, as assessed with the INCell high-content microscope using n= 3 independent wells per experimental condition.

**Supplementary figure 10.**
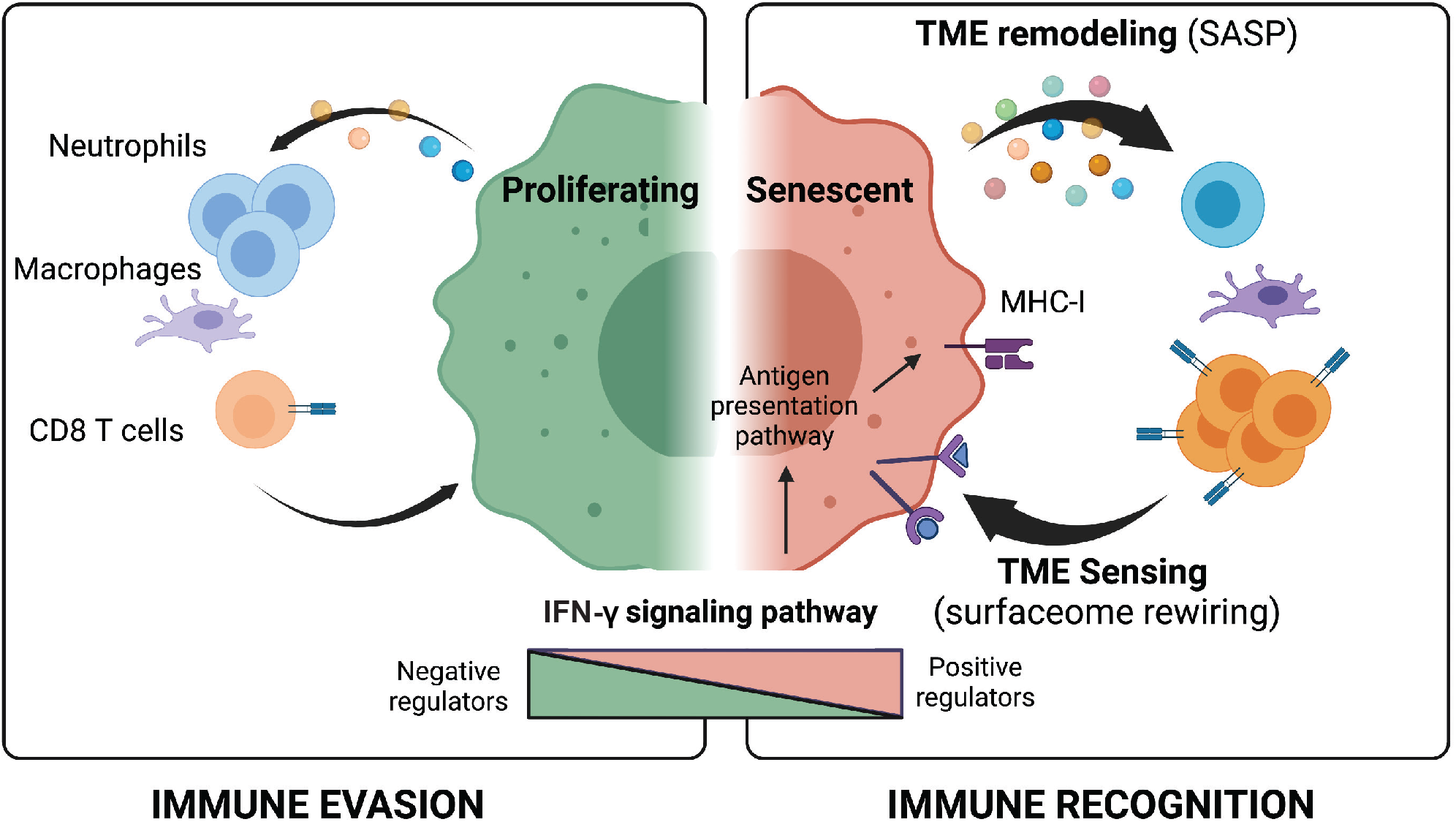
Graphic illustration of our working model.

## SUPPLEMENTARY TABLE

**Supplementary Table 1:** RNA-Seq data of proliferating (PRO) or senescent (SEN) NSP liver tumor cells, for both p53-restoration and drug-induced (trametinig+Palbociclib) settings. PRO and SEN cells were also treated with the BET inhibitor JQ-1 (500 n, 48 h), to expose BRD4-mediated transcriptional output in each cellular state.

## MATERIALS AND METHODS

### Cell Culture and drug treatment

p53-restorable mouse liver cancer cell lines were cultured in DMEM supplemented with 10% FBS and 1% penicillin and streptomycin (GIBCO) on plates that were collagen-coated (PurCol, Advanced Biomatrix, 0.1 mg/ml) for 30’ at 37 C and maintained by the addition of 1 μg/ml doxycycline to suppress p53 expression. In order to restore p53 expression and therefore induce senescence, doxycycline-containing media was replaced with doxycycline-free media for 6 to 8 days. Several cell lines have been generated and NSP is predominantly used for the study given the robustness of senescence phenotype upon p53 restoration. For human liver cell lines, HepG2 and SK-Hep1 were cultured with EMEM and SNU447 was cultured in RPMI-1640 in non-coated, tissue culture treated plates, all supplemented with 10% FBS and 1% penicillin and streptomycin. The concentration and regimen of drug treatment in cancer cell lines were as followed. For perturbing BRD4-dependent transcriptional programs, cells were treated with 500 nM of JQ-1 (S7110, Selleck Chem) for 48 h prior to harvest, starting JQ-1 at day 6 after restoring p53 (off-dox), when NSP cells are fully senescent. For drug-induced senescence experiments, p53-suppressed (on-dox) NSP cells were treated with trametinib (25 nM, S2673 Selleck Chem) + Palbociclib (500 nM, S1116, Selleck Chem), Nutlin (10 μM, S1061, Selleck Chem) or Cisplatin (1 μM), changed every 2-3 days, during 7 days. The concentration of DMSO corresponded to the drug treatment and does not exceed 1:1,000 dilution of total media volume, which shows no discernable toxicity to cultured cells. For IFN-γ of proliferating or senescent populations, the indicated doses of mouse or human recombinant IFN-γ was administrated to murine and human cancer cell lines respectively after 24 h of cell seeding and cells were harvested after 24 h of IFN-γ treatment for phenotypic or molecular analyses.

### Primary liver tumor generation and isolation of liver cell lines

C57BL/6N female mice aged 8-9 weeks old were injected via hydrodynamic tail vein injection (HTVI) with a sterile 2 ml (or 1/10 of mouse body weight) 0.9% NaCl solution containing 5 μg of pT3-EF1a-NrasG12D-IRES-rtTA (Tet-On system) and 20 ug of pT3-TRE-tRFP-shp53 transposon vectors along with 5ug CMV-SB13 transposase (5:1 ratio) through the lateral tail vein. Doxycycline was administered to mice via 625 mg/kg doxycycline-containing food pellets (Harlan Teklad) at least 4 days before injection. The tumor was harvested at 5-7 weeks after injection for cell line isolation. To derive cancer cell lines from primary liver tumor, tumors were minced and digested with 5ml of digesting solution, containing 1 mg/ml collagenase IV (C5138, Sigma-Aldrich) and 0.3% Dispase II (Roche 04942078001) in DMEM, at 37 °C for 30 mins with occasional vortexing. The cells were spun down to remove the supernatant and plated on collagen-coated plate. Independent cell lines were passaged at least 7-8 passages to remove fibroblasts and obtain homogenous population. For those experiments involving bioluminescence tracking of tumor growth the transposon construct pT3-EF1a-NrasG12D-IRES-rtTA-IRES-Luc was used. In the Tet-OFF system setting, the transposon construct pT3-EF1a-NrasG12D-IRES-tTA was used to co-inject with pT3-TRE-tRFP-shp53 vector into mice under normal diet to allow p53 hairpin expression. To restore p53 in the liver tumor, the mice were subjected to doxycycline diet. For constitutive p53 knockdown model, transposon constructs pT3-EF1a-NrasG12D and pT3-EF1a-tRFP-shp53 were used.

### Orthotopic transplant experiments

Both C57BL/6 mice were predominantly used for the animal study for the HTVI tumor generation and orthotopic liver injection experiments in the immunocompetent setting. C57BL/6N strain was mainly used except for the matching control strain with IFNG KO mice (Jax, #002287) that was in the C57BL/6J background. No difference was observed in terms of tumor growth or senescence surveillance phenotype between C57BL/6N and J strain. Female mice were used in the experiment for the convenience of cage separation. All in vivo experiments were performed with age-matched (8-13 weeks old) cohorts. For the orthotopic liver tumor injection, NSP tumor cells were trypsinized and filtered twice using 40 μm strainer to reduce cell doublets followed by pelleting and were prepared in 20 ul of 1:1 DMEM to Matrigel ratio and injected using 31-gauge needle to the left lobe of the mouse liver following the standard microsurgery institutional practice. Due to the engraftment differences in mice of different strains-C57BL/6, Nude and R2G2 (Envigo) mice-different amounts of tumor cells were injected. Specifically, 5x10^5^, 8x10^4^ and 5x10^4^ cells were injected respectively in each strain to have comparable tumor size around 2 weeks after injection. Mice were then randomized based on the similar size of tumor and assigned to different groups for the subsequent experimental design.

### Lentiviral and retroviral production and transduction

Lentiviruses were generated by co-transfection of viral vectors (1.5 μg) with packaging plasmids psPAX2 (0.75 μg) and pCMV-VSVG (0.25 μg) (Addgene) into 293T cells with 90% confluency in a 6-well plate. Retroviruses were generated by co-transfection of viral vectors (2 μg) with pCMV-VSVG (0.25 μg) (Addgene) into Phoenix-gp cells with 90% confluency in a 6-well plate. Polyethylenimine (PEI) was added during co-transfection with a ratio of total DNA:PEI = 1:3 to facilitate the binding of the plasmid to the cell surface. Viral containing supernatants were cleared of cellular debris by 0.45 μm filtration. Target cells were exposed to viral supernatants and mixed with 4 μg/ml polybrene for overnight before being washed, grown for 24 h in fresh media, then subjected to antibiotic selection or fluorescence-based cell sorting.

### Lentiviral and retroviral vectors

Murine liver cancer cells were infected with retroviral vector MSCV-Luc2-IRES-GFP (52) to enable bioluminescence imaging. For visualization and staining of liver tumor cells in vivo, tumor cells were infected with either the following lentiviral vectors specified in the figure legends, pRRL-<u>S</u>FFV-<u>G</u>FP-mir<u>E</u>(shRen)-PGK-<u>p</u>uromycin (SGEP was a gift from Johannes Zuber, Addgene #111170) or pRRL-EFS-GFP-shRen (generated through replacing SFFV with EFS promoter and removing antibiotic selection marker puromycin), to label the cells with GFP. For visualization of IFN-γ sensing, tumor cells were infected with the lentiviral IGS reporter construct described below.

### Genetic manipulation of cell line using CRISPR/Cas9

In order to knock out specific genes in mouse and human liver tumor cell lines, the plasmid pSpCas9(BB)-2A-GFP (PX458) (PX458 was a gift from Feng Zhang, Addgene #48138) in which a sgRNA targeting either an intergenic region of chromosome 8 (Ctrl) or the specific gene of interest was cloned. Cells were transiently transfected by PEI (2 μg plasmid and 6 ul PEI in 6 well plate with 60% confluency). Transfected cells were subsequently FACS sorted by GFP positivity 36-48 h post-transfection. For Ifngr1 and B2m KO experiment, PX458 transfected cells were first stained with IFNGR1 (2E2, biotin) followed by Streptavidin-APC staining, and MHC-I (H-2k^b^; AF6-88.5.5.3, ) antibody respectively and negative cells were sorted. Sorted population were further tested with IFN-γ to evaluate KO efficiency by using MHC-I induction as a proxy. In order to generate p53 KO human tumor cells, cells were electroporated following manufacturer’s instructions. Briefly, cells were trypsinized, washed in PBS once, and counted and then resuspended in Neon Buffer R. In parallel, 1 μg of Cas9 (ThermoFisher) and 1 μg of sgRNA were complexed for 15 min at room temperature to form the Cas9 RNP complex, which was then mixed with the cell aliquot. The cell/RNP mixture was electroporated (1400 V pulse voltage, 20 ms pulse width, 2 pulses) using Neon electroporation system (Thermo Fisher). The cells were recovered for 3 days with further selection through nutlin treatment (10 uM, Selleck Chemicals S1061) for 5-7 days to enrich p53 KO cells. The sgRNA sequence used in the experiments are: Ifngr1: TGGAGCTTTGACGAGCACTG, B2m: AGTATACTCACGCCACCCAC, Ctrl: GACATTTCTTTCCCCACTGG and TP53: CGCTATCTGAGCAGCGCTCA.

### Co-culture assays

In order to isolate CD8^+^ T cells from spleens of female OT-I mice (Jackson laboratory), spleens were mechanically disrupted by passing them through a 70 μm cell strainer and centrifuged at 1500 rpm x 5 minutes. Red blood cells were lysed with ACK lysis buffer (Quality Biological) for 5 minutes. Total splenocytes or CD8+ T cells FACS sorted on a Sony MA900 were then activated with CD3/CD28 Dynabeads (one bead/T cell, Thermo Fisher) and cultured in presence of IL-2 (2 ng/ml; Biolegend), IL-7 (2.5 ng/ml; Peprotech), IL-15 (50 ng/ml; Peprotech) and 2-mercaptoethanol (5.5uM, Fisher Scientific) in complete RPMI-1640 media supplemented with 10% FBS and 100 IU/ml penicillin/streptomycin for 5-6 days (passage cells every 2-3 days) prior to co-culture assays with mouse liver tumor cells. For Kupffer cells isolation, BL/6 male mice aged 8-14 weeks were first subjected to liver perfusion as previously described (53). After perfusion, the liver was removed and homogenized and then digested with protease solution (0.5 mg/ml type XIV protease, Sigma, P5147) supplemented with DNAseI (0.2 ug/ml, Roche, 10104159001) for 15 minutes at 37C with constant stirring. This suspension was then centrifuged at 50 g for 3 minutes to remove the hepatocyte pellet. The supernatant was then transferred and centrifuged 580 g for 5 minutes at 4C. Next, the pellet was washed with HBSS to remove residual protease solution and centrifuged at 580 g for 5 minutes at 4C to pellet the cells again. The pellet was then resuspended with FACS buffer and subjected to α-F4/80 isolation according to the manufacturer instruction (Miltenyi Biotec, 130-110-443). After isolation, the purity of Kupffer cells was confirmed with F4/80 staining through flow cytometry.

Murine liver tumor cells NSP were transduced with retrovirus expressing PresentER-SIINFEKL construct (GFP) (PresentER-SIINFEKL (GFP) was a gift from David Scheinberg, Addgene #102944) to express the peptide 257-264 from chicken ovalbumin, which is presented by H-2Kb on the cell surface. Transduced cells were further selected with puromycin to obtain > 95% GFP positivity. Tumor cells were cultured in presence or absence of doxycycline for 6 days in order to induce senescence in those cells were doxycycline was withdrawn. 1,000 proliferating or 2,000 senescent tumor cells were plated in the individual well of a 96 well collagen-coated plate. For those experiments where Kupffer cells were added, they were isolated on the same day and plated at the indicated ratio 6 h after plating the tumor cells. 24 h after plating tumor cells, previously activated OT-I T cells were added at the indicated ratio. Co-cultures were imaged over time using an INCell 6000 high-content imager (GE Healthcare Life Sciences), with a 488 nm and a 633 nm laser excitation to visualize tumor cells and T cells (stained by CellTracker Deep Red Dye, Invitrogen C34565) respectively, using a 10x objective. Images were captured at indicated time points, starting after the seeding of T cells onto tumor cells/Kupffer cells co-cultures. Images for each channel were saved during the experiment and subsequently analyzed using Columbus image analysis software. GFP+ tumor cells were identified and segmented from background using an intensity-based threshold method. T cells were identified using the same threshold method as the tumor cells. Number of the GFP+ tumor cells was quantified and normalized to the untreated control to calculate the killing index.

### Senescence *in vitro* and *in vivo* assays

For colony formation assays, 2,500 mouse liver cancer cells or 10,000 human liver cancer cells were plated in each well of a 6-well plate. Cells were cultured for 6 days, then fixed with 4% formaldehyde, and stained with crystal violet. Detection of SA-β-gal activity was performed as previously described at pH 5.5 for mouse cells and tissue and pH 6 for human cells (20). For in vivo SA-β-gal staining, fresh frozen tissue sections were fixed with 0.5% glutaraldehyde followed by standard SA-β-gal staining as above described. Sections were counterstained with eosin. For population doubling curves, cells were washed with PBS, trypsinized, and 100,000 cells were plated in triplicates in 6-well plates in presence or absence of doxycycline. Every 48 h cells were counted and 1 x 10^5^ cells were replated. Population doublings for each 48 h period were calculated by dividing the final cell number to initial cell number.

### Whole mount immunostaining and tissue clearing

To detect T cells and neutrophils in the NSP liver tumors, we performed whole mount immunostaining and tissue clearing (with benzyl alcohol, benzyl benzoate, <u>BABB)</u> of excised tumors as previously described (31). At the indicated time points, mice were euthanized by carbon dioxide inhalation and liver tumors collected and fixed in 4% paraformaldehyde in PBS at 4°C overnight. Tissues were washed three times with PBS for 10’ at room temperature and preserved in 0.05% azide in PBS at 4 °C before processing. Then, the tissues were permeabilized in methanol (MetOH) gradients in PBS (PBS > 50% MetOH > 80% MetOH > 100%MetOH, 30 min in each solution), bleached with Dent’s bleach (15% H_2_O_2_, 16.7% dimethyl sulfoxide [DMSO] in MetOH) for 1h at room temperature, and rehydrated through descending MetOH gradients in PBS (80% MetOH > 50% MetOH > PBS, 30 min in each solution). Tissues were next incubated in blocking buffer (0.3% Triton X100, 0.2% BSA, 5% DMSO, 0.1% azide and 25% FBS in PBS) for 24h at 4°C on a shaker and then stained with antibodies (rat anti-CD3 [clone 17A2, cat#100202, Biolegend]; goat anti-myeloperoxidase [goatMPO, AF3667, R&D Systems], and [hamsteranti-CD31, 2H8, MA3105, Thermo Fisher] all diluted 1:200 in blocking buffer), for 3 days at 4°C on a shaker. Tissues were next washed for 24 h in washing buffer (PBS with 0.2% Triton X100 and 3% NaCl), and stained with secondary antibodies (donkey anti-rat-AF488 [A212008, Invitrogen] and donkey anti-goat AF647 [A21447, Invitrogen] diluted at 1:400 in blocking buffer) for 2 days at 4°C with shaking. Tissues were then washed for 24 h in washing buffer and thereafter stained with goat anti-hamster-AF568 (goat anti-hamster IgG (H+L) cross-adsorbed secondary antibody, Alexa Fluor 568, A21112, Thermo Fisher, diluted at 1:400) and (1:1000) in blocking buffer for 2 days at 4°C, on a shaker. Tissues were then washed for 24 h in washing buffer and thereafter dehydrated in MetOH gradients in dH20 using glass containers (50% MetOH > 70% MetOH > 90% MetOH > 3x 100% MetOH, 30 min for each step). Tissues were next cleared for 30 min in 50% MetOH and 50% BABB (benzyl alcohol, benzyl benzoate, mixed 1:2) followed by clearing 1 h in 100% BABB. Finally, the tissues were imaged on an SP8 Microscope (Leica). Visualization and quantification was performed with Imaris software (Bitplane). In separate experiments, 3D imaging after tissue clearing was used to detect the ZsGreen1, IFN-γ sensing (IGS) reporter. For these experiments, we used the <u>CUBIC</u> tissue clearing protocol that maintains the fluorescence from fluorescent proteins (54). Tissues were excised and fixed as stated above, and then were soaked in CUBIC-I solution in a 15 mL conical tube container. CUBIC-I was prepared mixing 108 ml of ddH2O with 75g of Urea (Sigma, U5128), 75g of N,N,N’,N’-Tetrakis(2-Hydroxypropyl)ethylenediamine (Sigma, 122262) and 42ml of Triton X-100 (Sigma, X100). Samples were maintained at 37°C on a shaker for 7 days, changing the media every other day, until clear. The samples were then counterstained for DAPI in CUBIC-1 (1:1000) for 24h and washed in CUBIC-I overnight. Images were acquired and analyzed as described above.

### Western blotting

Cell were lysed with RIPA buffer (50 m Tris PH 7.4, 150 mM NaCl, 0.5 % sodium deoxycholate, 0.1% SDS; 1mM EDTA; 1% NP-40) supplemented with phosphatase and protease inhibitor (5872, Cell Signaling Technology) and protein concentration was determined by BCA assay. Samples were boiled for 5 minutes and 20 to 30 μg of protein were separated by SDS-PAGE, transferred to polyvinylidene difluoride (PVDF) membranes (Millipore) according to standard protocols and probed with the relevant primary antibody overnight at 4°C. Membranes were then incubated with horseradish peroxide (HRP)-conjugated) anti-rabbit IgG or anti-mouse IgG secondary antibodies (1:10,000, GE Healthcare Life Science) at room temperature and proteins were detected using Pierce ECL Western Blotting Substrate (34095, Thermo Fisher Scientific). Antibodies were diluted as follows: p53 (CM5) (1:500, NCL-L-p53-CM5p, Leica Biosystems), p21 (F-5) (1:500, sc-6246, Santa Cruz Biotechnology), Phospho-Stat1 (Tyr701) (1:500, #9167, Cell Signaling Technology), Stat1 (1:1,000, #14994, Cell Signaling Technology), TCPTP (Ptpn2, 1:1000, ab180764, Abcam). Protein loading was measured using a monoclonal β-actin antibody directly conjugated to horseradish peroxidase (1:20,000; AC-15, Sigma-Aldrich), nucleolin (1:5000, ab22758, Abcam) or vinculin (1:2,000, ab129002, Abcam). ECL developed blots were imaged using a FluorChem M system (Protein Simple).

### In vitro multiplexed ELISA

Conditioned media samples (duplicates collected in complete DMEM 48 h after seeding) from proliferating or senescent NSP tumor cells (6 to 8 days after doxycycline withdrawal) were centrifuged at 1500 rpm for 3 minutes and filtered through 0.2 um filter to remove cell debris. Samples concentrations were normalized by diluting in complete DMEM according to cell count. Aliquots (50 μl) of the conditioned media were analyzed using multiplex immunoassays designed for mouse (Mouse Cytokine/Chemokine Array 31-Plex) from Eve Technologies. Biological replicates from two independent experiments were performed to determine cytokine levels. Heatmaps display relative cytokine expression values normalized to geometric means of individual cytokines from both proliferating and senescent samples.

### Measurement of IFN-γ in *in vivo* tumor lysates

BD cytometric bead array Mouse Th1/Th2 cytokine kit (Cat# 551287, BD Biosciences) was used to determine the IFN-γ levels. Flash frozen tissues were lysed in RIPA buffer and homogenized using TissueLyser II (Qiagen) followed by protein concentration measurement determined by BCA assay. 100 ug of tissue lysate were used subsequent measurement following standard manufacturer instructions of CBA kits.

### Plasma membrane-enriched mass spectrometry

To capture differential cell surface proteome changes induced by senescence, we adapted the protocol from previous published study (38) and followed the manufacturer instruction (Pierce Cell Surface Protein Isolation Kit #89881) to enrich cell surface proteins of proliferating and senescent cells through biotin-based labeling followed by pull-down purification. In brief, we plated 1 and 3 15 cm plates of proliferating and senescent cells (6 days after doxycycline withdrawal) with an initial seeding of 7x10^5^ and 2x10^6^ million cells respectively and collected the cells 2 days later, with the cells approximatively at 85% confluency. Before harvesting the cells, cells were incubated with biotin solution for 30 minutes at 4C to allow the surface protein labeling. Cells were then washed with cold PBS and scraped down followed by lysis (buffer provided in the kit). Lysates were centrifuged and the clarified supernatant was used for purification of biotinylated proteins on NeutrAvidin Agarose. Supernatant was incubated with NeutrAvidin Agarose for 2 h at room temperature in the closed column to allow biotinylated proteins binding. Column containing Agarose slurry was washed to remove unbound proteins. The proteins were then digested in situ in the column overnight using 4μg of trypsin (Promega, V5111) per column at 37C on a rotor. Digested proteins were further desalted by C18 Stagetip and subjected to liquid chromatography– mass spectrometry (LC-MS/MS) followed by proteins identification through Proteome Discover (Thermo Scientific) according to protocols previously described (38). Non-biotinylated cell lysates were also included and served as background controls.

### Protein identification

The LC-MS/MS .raw files were processed using Mascot and searched for protein identification against the SwissProt protein database for human/mouse (please adjust the species accordingly). Carbamidomethylation of C was set as a fixed modification and the following variable modifications allowed: oxidation (M), N-terminal protein acetylation, deamidation (N and Q), and phosphorylation (S, T and Y). Search parameters specified an MS tolerance of 10 ppm, an MS/MS tolerance at 0.080 Da and full trypsin digestion, allowing for up to two missed cleavages. False discovery rate was restricted to 1% in both protein and peptide level. Normalized protein intensities were obtained using Scaffold (4.8.4).

### RNA preparation and High throughput RNA-sequencing analysis

For in vitro liver cell lines RNA preparation, total RNA was extracted using using TRIzol (Thermo Fisher Scientific) following the manufacturer’s instructions. For in vivo bulk tumor RNA-seq, proliferating tumor (p53 Off) was harvested 7-10 day after randomization point and senescent-induced tumor (p53 On) was harvested 12 days after p53 restoration, allowing similar size of tumor at harvest. To extract tissue RNA, freshly isolated tumor chunk was first stored in RNA-later solution (AM7024, Thermo Scientific) to preserve RNA integrity until extraction and RNeasy kit (74106, Quiagen) was used to purified tissue RNA following the manufacturer instructions. Purified polyA mRNA was subsequently fragmented, and first and second strand cDNA synthesis performed using standard Illumina mRNA TruSeq library preparation protocols. Double stranded cDNA was subsequently processed for TruSeq dual-index Illumina library generation. For sequencing, pooled multiplexed libraries were run on a HiSeq 2500 machine on RAPID mode. Approximately 10 million 76bp single-end reads were retrieved per replicate condition. Resulting RNA-Seq data was analyzed by removing adaptor sequences using Trimmomatic (55), aligning sequencing data to GRCm38 – mm10 with STAR (56), and genome wide transcript count was quantified using featureCounts (57) to generate raw count matrix. Differential gene expression analysis was performed using DESeq2 package (58) between experimental conditions, using 3 independent biological replicates (independent cultures of NSP tumor cells) per condition, implemented in R (http://cran.r-project.org/). Differentially expressed genes (DEGs) were determined by > 2-fold change in gene expression with adjusted P-value < 0.05. For heatmap visualization of DEGs, samples were z-score normalized and plotted using ‘pheatmap’ package in R. Functional enrichments of these differential expressed genes were performed with enrichment analysis tool Enrichr (59). Gene expressions of RNA-Seq data were clustered using hierarchical clustering based on one minus Pearson correlation test. Subtype specific gene signatures were derived (25) and used as inputs for signature score calculation using R package singscore (60).

### Public dataset transcriptomic analyses

Signature of different human liver cancer subtype was obtained from previous study (25). In brief, the top 200 over-expressed and under-expressed gene transcripts among each tumor subtype were selected as their signature. To analyze the transcriptomic changes of genes encoding plasma membrane and extracellular factors distinguishing senescent and proliferating tumor cells, transcriptomic data of a series of human tumor cell lines triggered to senesce was used according to the previously published study (37) and obtained from the website https://ccb.nki.nl/publications/cancer-senescence/. The expression of selected genes was compared between senescent and the corresponding proliferating cells among individual cell lines and normalized to determine the fold change. Information about protein subcellular localization was derived from the Compartments_knowledge_based database (61), with the genes assigned to specific subcellular localization when the criteria score is >= 3. The Cancer Genome Atlas Liver Hepatocellular Carcinoma (TCGA-LIHC) data set, including p53 mutational status, transcriptomic profiles, and patient survival, were downloaded using R package TCGAbiolinks (62, 63). Senescence signatures derived from our mouse models were used as input for computing signature scores using ssgsea method in R package GSVA (64). These signature scores were used to separate patients into high and low groups, and log rank test was used to test the differences in survival between these two groups.

### Gene set enrichment analysis (GSEA)

GSEA was performed using the GSEAPreranked tool for conducting gene set enrichment analysis of data derived from RNA-seq experiments (version 2.07) against signatures in the MSigDB database (http://software.broadinstitute.org/gsea/msigdb), signatures derived herein, and published expression signatures in organoid models and human samples. The metric scores were calculated using the sign of the fold change multiplied by the inverse of the p-value.

### Reverse transcription and quantitative PCR

Total RNA was isolated from mouse liver tumor cell line using TRIzol (Thermo Fisher Scientific) following the manufacturer’s instructions. cDNA was obtained from 500 ng RNA using the Transcriptor First Strand cDNA Synthesis Kit (04896866001, Roche) after treatment with DNaseI (18068015, Thermo Fisher Scientific) following the manufacturer’s instructions using random hexamer method. The following primer sets for mouse sequences were used: Tap1_F 5’-GGACTTGCCTTGTTCCGAGAG-3’, Tap1_R 5’-GCTGCCACATAACTGATAGCGA-3’, Psmb8_F 5’-ATGGCGTTACTGGATCTGTGC-3’, Psmb8_R 5’-CGCGGAGAAACTGTAGTGTCC-3’, Nlrc5_F 5’-CCTGCGTCCCAGTCATTC-3’, Nlrc5_R 5’-CTGCTGGTCAGTGATGGAGA-3’, Erap1_F 5’-TAATGGAGACTCATTCCCTTGGA-3’, Erap1_R 5’-AAAGTCAGAGTGCTGAGGTTT G-3’, H2-K1_F 5’-GCTGGTGAAGCAGAGAGACTCAG-3’, H2-K1_R 5’-GGTGACTTTATCTTCAGGTCTGCT-3’, H2-D1_F 5’-AGTGGTGCTGCAGAGCATTACAA-3’, H2-D1_R 5’-GGTGAC TTCACCTTTAGATCTGGG-3’, B2m_F 5’-TTCTGGTGCTTGTCTCACTGA-3’, B2m_R 5’-CAG TATGTTCGGCTTCCCATTC-3’, Hprt_F 5′-TCAGTCAACGGGGGACATAAA-3′, Hprt_R 5′-GGGGCTGTACTGCTTAACCAG-3′, Rplp0_F 5′-GCTCCAAGCAGATGCAGCA-3′, Rplp0_R 5′-CCGGATGTGAGGCAGCAG-3′, Quantitative PCR with reverse transcription (qRT–PCR) was carried out in triplicate (10 cDNA ng per reaction) using SYBR Green PCR Master Mix (Applied Biosystems) on the ViiA 7 Real-Time PCR System (Life technologies). Hprt, Rplp0 (also known as 36b4) served as endogenous normalization controls.

### Tumor measurement by ultrasound and bioluminescence imaging

High-contrast ultrasound imaging was performed on a Vevo 2100 System with a MS250 13-to 24-MHz scanhead (VisualSonics) to stage and quantify liver tumor burden. Tumor volume was analyzed using Vevo LAB software. Bioluminescence imaging was used to track luciferase expression in orthotopically injected liver tumor cells expressing a Luc-GFP reporter as well as primary HTVI tumor harboring luciferase construct (vector described above). Mice were injected IP with luciferin (5 mg/mouse; Gold Technologies) and then imaged on a Xenogen IVIS Spectrum imager (PerkinElmer) 10 minutes later. Quantification of luciferase signaling was analyzed using Living Image software (Caliper Life Sciences).

### Flow cytometry and sample preparation

For *in vivo* sample preparation, orthotopically injected liver tumors were isolated by removing the adjacent normal tissue, and allocated for 10% formalin fixation, OCT frozen blocks, snap frozen tissue, and flow cytometry analysis. To prepare single cell suspensions for flow cytometry analysis, liver tumor was mechanically disrupted to a single cell suspension using a 150 μm metal mesh and glass pestle in ice-cold 3% FBS/HBSS and passed through a 70 μm strainer. The liver homogenate was spun down at 400 g for 5 minutes at 4°C, and the pellet was resuspended in 15ml 3% FCS/HBSS, 500ul (500U) heparin, and 8ml Percoll (GE), mixed by inversion, and spun at 500 g for 10 min at 4°C. After removal of supernatant, cells were resuspended in PBS supplemented with 2% FBS. Samples were blocked with anti-CD16/32 (1:200, FC block, #553142) (BD Pharmigen) for 20 minutes and then incubated with the following antibodies for 30 minutes on ice: CD3 (1:200, 17A2, #612803), CD19 (1:200, 1D3, #563235), CD4 (1:800, RM4-5, #563151),

CD44 (1:200, IM7, #560568), CD11b (1:800, M1/70, #563553) (BD Biosciences); MHC-I (1:100, H-2k^b^; AF6-88.5.5.3, #17-5958-82), CD119 (1:100, 2E2, #13-1191-82), Armenian Hamster IgG isotype (1:100, eBio299Arm, #13488881) (Invitrogen); CD45 (1:400, 30-F11M, #103128), Gr-1 (1:200, RB6-8C5, #108406), F4/80 (1:100, BM8, #123116), CD8 (1:400, 53-6.7, #100721), Ly6C (1:200, HK1.4, # 128026), CD11c (1:200, N418, #117335), Ly6G (1:200, 1A8, #563005), CD69 (1:200, H1.2F3, #104522), CD106 (1:100, MVCAM.A, #105717), CD62L (1:200, MEL-14, #104435), PD-1 (1:100, 29F.1A12, #135215) (Biolegend); Streptavidin (1:200, #20-4317-U100), TIGIT (1:100, 1G9, #20-1421-U025), NK1.1 (1:100, PK136, #65-5941-U100) (Tonbo); human antibody HLA-A,B,C (1:100, W6/32, #17-9983-42) (Thermo Fisher). To distinguish live/dead cells, DAPI and Ghost dye violet 510 (1:1000, #13-0870-T100) (Tonbo) were used depending on whether the cells are fixed. For fixed cells, cells were stained in PBS prior to antibody staining. Flow cytometry was performed on an LSRFortessa or Guava flow cytometer (Luminex Corporation), and data were analyzed using FlowJo (TreeStar).

### Neutralizing antibody and liposomal clodronate studies

To determine the specific immune cell dependency of senescence surveillance, depleting antibodies or drugs were administrated to the mice one day after doxycycline withdrawal. For NK cell depletion, mice were injected intraperitoneally (IP) with an α-NK1.1 antibody (250 μg; PK136, BioXcell) twice per week. For T cell depletion, mice were injected IP with either an α-CD4 (200 μg; GK1.5, BioXcell) or α-CD8 antibody (200 μg; 2.43, BioXcell) twice per week. Depletion of NK, CD4+, and CD8+ T cells was confirmed by flow cytometric analysis of liver tumor tissue. For neutrophil/myeloid-derived suppressive cells depletion, mice were injected intraperitoneally with an α-Gr-1 (200 μg; RB6-8C5, BioXcell) twice per week. For control, isotype control antibody (200 μg; LTF-2, BioXcell) was IP twice per week. For macrophage depletion, mice were injected intravenously (IV) with clodronate liposomes (10 gram/kg of mouse weight; ClodronateLiposomes.com) twice per week. PBS was used as a control.

### Immunofluorescence and immunohistochemistry

Tissues were fixed overnight in 10% neutral buffered formalin (Richard-Allan Scientific), embedded in paraffin and cut into 5 µm sections. Sections were deparaffinized and rehydrated with a histoclear/alcohol series and subjected to antigen retrieval by boiling in citrate antigen retrieval buffer (Vector). Slides were then blocked in PBS/0.1% Triton X-100 containing 1% BSA. Primary antibodies were incubated overnight at 4°C in blocking buffer. The following primary antibodies were used: GFP (ab13970, Abcam, 1:500), Ki67 (#550609, BD Biosciences, 1:200), CD8 (4SM15, eBioscience, 1:200), CD45 (#70257, Cell Signaling Technology, 1:100), F4/80 (#70076, Cell Signaling Technology, 1:200). For immunohistochemistry, Vector ImmPress HRP kits and ImmPact DAB (Vector Laboratories) were used for secondary detection. For immunofluorescence, the following secondary antibodies were used: goat anti-chicken AF488 (A11039, Invitrogen, 1:500), donkey anti-rabbit AF594 (A21207, Invitrogen, 1:500), goat anti-rabbit AF594 (A11037, Invitrogen, 1:500), donkey anti-rabbit AF647 (A31573, Invitrogen, 1:500). All secondary antibodies were diluted in blocking buffer and incubated for 1 h at room temperature. Subsequently, slides were washed and nuclei were counterstained with PBS containing DAPI (1 μg/ml), and mounted under cover slips with ProLong Gold (Life Technologies). Images were acquired with a Zeiss AxioImager microscope using Axiovision software.

### Generation of IFN-γ sensing (IGS) reporter

In order to generate the IFN-γ sensing reporter from our study, we have adapted the construct design from the previously described paper (50). In brief, we have crafted a 5x Interferon Gamma-activated sequence (GAS) inserted in front of a mini promoter (minimal TATA-box promoter with low basal activity) followed by ZsGreen1 reporter. Right after the reporter sequence, this lentiviral construct also contains RFP driven by the PGK promoter to have constitutive RFP expression for cell visualization. The cells were transduced with virus and sorted through flow cytometry with high RFP level for stable expression of the construct in the cells.

### Statistical analyses

Statistical analyses were performed as described in the figure legend for each experiment. Group size was determined on the basis of the results of preliminary experiments, and no statistical method was used to predetermine sample size. The indicated sample size (n) represents biological replicates. All samples that met proper experimental conditions were included in the analysis. Particularly, we have observed that in the orthotopic transplantation setting, the undesired lung metastasis (lung weight > 300 mg) occurred due to the technical limitation of liver injection may affect the tumor regression phenotype upon p53 restoration and the mice were thus excluded for the analysis. Survival was measured using the Kaplan–Meier method. Statistical significance was determined by Student t test, log-rank test, Mann–Whitney test, Fisher exact test, and Pearson correlation using Prism 6 Software (GraphPad Software) as indicated. Significance was set at P < 0.05.

### Figure Preparation

Figures were prepared using BioRender.com for scientific illustrations and Illustrator CC 2020 (Adobe).

### Data Availability

RNA-seq data generated in this study are deposited in the Gene Expression Omnibus database under accession number GSE203140.

## Authors’ Contributions

**Conceptualization and design:** H.-A. Chen, D. Alonso-Curbelo, S.W. Lowe

**Methodology:** H.-A. Chen, R. Mezzadra, J.M. Adrover, C. Zhu, R.C. Hendrickson, D. Alonso-Curbelo

**Acquisition and analysis of data (e.g., investigation, validation, resources):** H.-A, Chen, R. Mezzadra, J.M. Adrover, C. Zhu, W. Luan, A. Wuest, S. Tian, X. Li, R.C. Hendrickson, M. Egeblad, D. Alonso-Curbelo

**Formal analysis and software:** Y.-J. Ho, J.M. Adrover, R.C. Hendrickson

**Writing–original draft presentation:** H.-A, Chen, D. Alonso-Curbelo, S.W. Lowe

**Writing–review and editing:** H.-A, Chen, Y.-J. Ho, R. Mezzadra, J.M. Adrover, C. Zhu, M. Egeblad, D. Alonso-Curbelo

**Visualization**: H.-A, Chen, Y.-J. Ho, .M. Adrover, D. Alonso-Curbelo

**Study supervision:** D. Alonso-Curbelo, S.W. Lowe

## Acknowledgments

We thank J. Simon and the MSKCC animal facility for technical support with animal colonies; J.E. Wilkinson and J. Shia for mouse liver cancer pathology consulting; E. de Stanchina and A. Kulick from the MSKCC Antitumor Assessment Core Facility for animal treatment support; Z. Li, A. Gaito and M. Miele from the MSKCC Microchemistry & Proteomics core for proteomic profiling technical support; and J.P. Morris IV, S. Houlihan and other members of the Lowe laboratory for advice and discussions. H.-A. Chen was supported by a NIH F99 Grant (F99CA245797). D. Alonso-Curbelo was supported by the the *La Caixa* Junior Leader Fellowship (LCF/BQ/PI20/11760006). R. Mezzadra was supported by NWO (Dutch Research Council-Rubicon Fellowship 452182318) and is a Cancer Research Institute Irvington Fellow supported by the Cancer Research Institute (CRI Award #3441). J.M. Adrover is the recipient of a Cancer Research Institute/Irvington Postdoctoral Fellowship (CRI Award #3435). C. Zhu is supported by an F32 Postdoctoral Fellowship (1F32CA257103) from the NIH/NCI. M. Egeblad is supported by a grant from the National Cancer Institute (1R01CA237413) and the work was supported by the Cold Spring Harbor Laboratory (CSHL) Cancer Center (P30CA045508). This work is also supported by P01CA087497 Roles and Regulation of wild-type and mutant p53 grant (to S.W. Lowe). S.W. Lowe is an investigator in the Howard Hughes Medical Institute and the Geoffrey Beene Chair for Cancer Biology.

## REFERENCE

1. Di Micco R, Krizhanovsky V, Baker D, d’Adda di Fagagna F. Cellular senescence in ageing: from mechanisms to therapeutic opportunities. Nature reviews Molecular cell biology 2021;22(2):75–95 doi 10.1038/s41580-020-00314-w.

2. Munoz-Espin D, Serrano M. Cellular senescence: from physiology to pathology. Nature reviews Molecular cell biology 2014;15(7):482–96 doi 10.1038/nrm3823.

3. Demaria M, Ohtani N, Youssef SA, Rodier F, Toussaint W, Mitchell JR, et al. An essential role for senescent cells in optimal wound healing through secretion of PDGF-AA. Developmental cell 2014;31(6):722–33 doi 10.1016/j.devcel.2014.11.012.

4. Xu M, Pirtskhalava T, Farr JN, Weigand BM, Palmer AK, Weivoda MM, et al. Senolytics improve physical function and increase lifespan in old age. Nat Med 2018;24(8):1246–56 doi 10.1038/s41591-018-0092-9.

5. Serrano M, Lin AW, McCurrach ME, Beach D, Lowe SW. Oncogenic ras provokes premature cell senescence associated with accumulation of p53 and p16INK4a. Cell 1997;88(5):593–602 doi 10.1016/s0092-8674(00)81902-9.

6. Ewald JA, Desotelle JA, Wilding G, Jarrard DF. Therapy-induced senescence in cancer. Journal of the National Cancer Institute 2010;102(20):1536–46 doi 10.1093/jnci/djq364.

7. Coppe JP, Desprez PY, Krtolica A, Campisi J. The senescence-associated secretory phenotype: the dark side of tumor suppression. Annual review of pathology 2010;5:99–118 doi 10.1146/annurev-pathol-121808-102144.

8. Demaria M, O’Leary MN, Chang J, Shao L, Liu S, Alimirah F, et al. Cellular Senescence Promotes Adverse Effects of Chemotherapy and Cancer Relapse. Cancer discovery 2017;7(2):165–76 doi 10.1158/2159-8290.CD-16-0241.

9. Coppe JP, Patil CK, Rodier F, Sun Y, Munoz DP, Goldstein J, et al. Senescence-associated secretory phenotypes reveal cell-nonautonomous functions of oncogenic RAS and the p53 tumor suppressor. PLoS biology 2008;6(12):2853–68 doi 10.1371/journal.pbio.0060301.

10. Tasdemir N, Banito A, Roe JS, Alonso-Curbelo D, Camiolo M, Tschaharganeh DF, et al. BRD4 Connects Enhancer Remodeling to Senescence Immune Surveillance. Cancer discovery 2016;6(6):612–29 doi 10.1158/2159-8290.CD-16-0217.

11. Chien Y, Scuoppo C, Wang X, Fang X, Balgley B, Bolden JE, et al. Control of the senescence-associated secretory phenotype by NF-kappaB promotes senescence and enhances chemosensitivity. Genes & development 2011;25(20):2125–36 doi 10.1101/gad.17276711.

12. Kuilman T, Michaloglou C, Vredeveld LC, Douma S, van Doorn R, Desmet CJ, et al. Oncogene-induced senescence relayed by an interleukin-dependent inflammatory network. Cell 2008;133(6):1019–31 doi 10.1016/j.cell.2008.03.039.

13. Basisty N, Kale A, Jeon OH, Kuehnemann C, Payne T, Rao C, et al. A proteomic atlas of senescence-associated secretomes for aging biomarker development. PLoS biology 2020;18(1):e3000599 doi 10.1371/journal.pbio.3000599.

14. Hernandez-Segura A, de Jong TV, Melov S, Guryev V, Campisi J, Demaria M. Unmasking Transcriptional Heterogeneity in Senescent Cells. Curr Biol 2017;27(17):2652–60 e4 doi 10.1016/j.cub.2017.07.033.

15. Xue W, Zender L, Miething C, Dickins RA, Hernando E, Krizhanovsky V, et al. Senescence and tumour clearance is triggered by p53 restoration in murine liver carcinomas. Nature 2007;445(7128):656–60 doi 10.1038/nature05529.

16. Kang TW, Yevsa T, Woller N, Hoenicke L, Wuestefeld T, Dauch D, et al. Senescence surveillance of pre-malignant hepatocytes limits liver cancer development. Nature 2011;479(7374):547–51 doi 10.1038/nature10599.

17. Ovadya Y, Landsberger T, Leins H, Vadai E, Gal H, Biran A, et al. Impaired immune surveillance accelerates accumulation of senescent cells and aging. Nature communications 2018;9(1):5435 doi 10.1038/s41467-018-07825-3.

18. Lujambio A. To clear, or not to clear (senescent cells)? That is the question. Bioessays 2016;38 Suppl 1:S56–64 doi 10.1002/bies.201670910.

19. Di Mitri D, Toso A, Chen JJ, Sarti M, Pinton S, Jost TR, et al. Tumour-infiltrating Gr-1+ myeloid cells antagonize senescence in cancer. Nature 2014;515(7525):134–7 doi 10.1038/nature13638.

20. Ruscetti M, Leibold J, Bott MJ, Fennell M, Kulick A, Salgado NR, et al. NK cell-mediated cytotoxicity contributes to tumor control by a cytostatic drug combination. Science 2018;362(6421):1416–22 doi 10.1126/science.aas9090.

21. Ruscetti M, Morris JPt, Mezzadra R, Russell J, Leibold J, Romesser PB, et al. Senescence-Induced Vascular Remodeling Creates Therapeutic Vulnerabilities in Pancreas Cancer. Cell 2020;181(2):424–41 e21 doi 10.1016/j.cell.2020.03.008.

22. Wagner V, Gil J. Senescence as a therapeutically relevant response to CDK4/6 inhibitors. Oncogene 2020;39(29):5165–76 doi 10.1038/s41388-020-1354-9.

23. Eggert T, Wolter K, Ji J, Ma C, Yevsa T, Klotz S, et al. Distinct Functions of Senescence-Associated Immune Responses in Liver Tumor Surveillance and Tumor Progression. Cancer cell 2016;30(4):533–47 doi 10.1016/j.ccell.2016.09.003.

24. Suda T, Liu D. Hydrodynamic gene delivery: its principles and applications. Mol Ther 2007;15(12):2063–9 doi 10.1038/sj.mt.6300314.

25. Chiang DY, Villanueva A, Hoshida Y, Peix J, Newell P, Minguez B, et al. Focal gains of VEGFA and molecular classification of hepatocellular carcinoma. Cancer research 2008;68(16):6779–88 doi 10.1158/0008-5472.CAN-08-0742.

26. Hoshida Y, Nijman SM, Kobayashi M, Chan JA, Brunet JP, Chiang DY, et al. Integrative transcriptome analysis reveals common molecular subclasses of human hepatocellular carcinoma. Cancer research 2009;69(18):7385–92 doi 10.1158/0008-5472.CAN-09-1089.

27. Llovet JM, Kelley RK, Villanueva A, Singal AG, Pikarsky E, Roayaie S, et al. Hepatocellular carcinoma. Nat Rev Dis Primers 2021;7(1):6 doi 10.1038/s41572-020-00240-3.

28. Simon S, Labarriere N. PD-1 expression on tumor-specific T cells: Friend or foe for immunotherapy? Oncoimmunology 2017;7(1):e1364828 doi 10.1080/2162402X.2017.1364828.

29. Kaech SM, Hemby S, Kersh E, Ahmed R. Molecular and functional profiling of memory CD8 T cell differentiation. Cell 2002;111(6):837–51 doi 10.1016/s0092-8674(02)01139-x.

30. Bruix J, Cheng AL, Meinhardt G, Nakajima K, De Sanctis Y, Llovet J. Prognostic factors and predictors of sorafenib benefit in patients with hepatocellular carcinoma: Analysis of two phase III studies. Journal of hepatology 2017;67(5):999–1008 doi 10.1016/j.jhep.2017.06.026.

31. Adrover JM, Aroca-Crevillen A, Crainiciuc G, Ostos F, Rojas-Vega Y, Rubio-Ponce A, et al. Programmed ’disarming’ of the neutrophil proteome reduces the magnitude of inflammation. Nature immunology 2020;21(2):135–44 doi 10.1038/s41590-019-0571-2.

32. Zeisberger SM, Odermatt B, Marty C, Zehnder-Fjallman AH, Ballmer-Hofer K, Schwendener RA. Clodronate-liposome-mediated depletion of tumour-associated macrophages: a new and highly effective antiangiogenic therapy approach. Br J Cancer 2006;95(3):272–81 doi 10.1038/sj.bjc.6603240.

33. De Simone G, Andreata F, Bleriot C, Fumagalli V, Laura C, Garcia-Manteiga JM, et al. Identification of a Kupffer cell subset capable of reverting the T cell dysfunction induced by hepatocellular priming. Immunity 2021;54(9):2089–100 e8 doi 10.1016/j.immuni.2021.05.005.

34. Milanovic M, Fan DNY, Belenki D, Dabritz JHM, Zhao Z, Yu Y, et al. Senescence-associated reprogramming promotes cancer stemness. Nature 2018;553(7686):96–100 doi 10.1038/nature25167.

35. Yousef H, Czupalla CJ, Lee D, Chen MB, Burke AN, Zera KA, et al. Aged blood impairs hippocampal neural precursor activity and activates microglia via brain endothelial cell VCAM1. Nat Med 2019;25(6):988–1000 doi 10.1038/s41591-019-0440-4.

36. Rapisarda V, Borghesan M, Miguela V, Encheva V, Snijders AP, Lujambio A, et al. Integrin Beta 3 Regulates Cellular Senescence by Activating the TGF-beta Pathway. Cell reports 2017;18(10):2480–93 doi 10.1016/j.celrep.2017.02.012.

37. Jochems F, Thijssen B, De Conti G, Jansen R, Pogacar Z, Groot K, et al. The Cancer SENESCopedia: A delineation of cancer cell senescence. Cell reports 2021;36(4):109441 doi 10.1016/j.celrep.2021.109441.

38. Perna F, Berman SH, Soni RK, Mansilla-Soto J, Eyquem J, Hamieh M, et al. Integrating Proteomics and Transcriptomics for Systematic Combinatorial Chimeric Antigen Receptor Therapy of AML. Cancer cell 2017;32(4):506–19 e5 doi 10.1016/j.ccell.2017.09.004.

39. Hoare M, Ito Y, Kang TW, Weekes MP, Matheson NJ, Patten DA, et al. NOTCH1 mediates a switch between two distinct secretomes during senescence. Nature cell biology 2016;18(9):979–92 doi 10.1038/ncb3397.

40. Castro F, Cardoso AP, Goncalves RM, Serre K, Oliveira MJ. Interferon-Gamma at the Crossroads of Tumor Immune Surveillance or Evasion. Frontiers in immunology 2018;9:847 doi 10.3389/fimmu.2018.00847.

41. Platanias LC. Mechanisms of type-I- and type-II-interferon-mediated signalling. Nature reviews Immunology 2005;5(5):375–86 doi 10.1038/nri1604.

42. Manguso RT, Pope HW, Zimmer MD, Brown FD, Yates KB, Miller BC, et al. In vivo CRISPR screening identifies Ptpn2 as a cancer immunotherapy target. Nature 2017;547 (7664):413ߝ8 doi 10.1038/nature23270.

43. Song MM, Shuai K. The suppressor of cytokine signaling (SOCS) 1 and SOCS3 but not SOCS2 proteins inhibit interferon-mediated antiviral and antiproliferative activities. J Biol Chem 1998;273(52):35056–62 doi 10.1074/jbc.273.52.35056.

44. Kleppe M, Soulier J, Asnafi V, Mentens N, Hornakova T, Knoops L, et al. PTPN2 negatively regulates oncogenic JAK1 in T-cell acute lymphoblastic leukemia. Blood 2011;117(26):7090–8 doi 10.1182/blood-2010-10-314286.

45. Zhou F. Molecular mechanisms of IFN-gamma to up-regulate MHC class I antigen processing and presentation. Int Rev Immunol 2009;28(3-4):239–60 doi 10.1080/08830180902978120.

46. McCarthy MK, Weinberg JB. The immunoproteasome and viral infection: a complex regulator of inflammation. Front Microbiol 2015;6:21 doi 10.3389/fmicb.2015.00021.

47. Yoshihama S, Vijayan S, Sidiq T, Kobayashi KS. NLRC5/CITA: A Key Player in Cancer Immune Surveillance. Trends in cancer 2017;3(1):28–38 doi 10.1016/j.trecan.2016.12.003.

48. Kalaora S, Lee JS, Barnea E, Levy R, Greenberg P, Alon M, et al. Immunoproteasome expression is associated with better prognosis and response to checkpoint therapies in melanoma. Nature communications 2020;11(1):896 doi 10.1038/s41467-020-14639-9.

49. Efeyan A, Ortega-Molina A, Velasco-Miguel S, Herranz D, Vassilev LT, Serrano M. Induction of p53-dependent senescence by the MDM2 antagonist nutlin-3a in mouse cells of fibroblast origin. Cancer research 2007;67(15):7350–7 doi 10.1158/0008-5472.CAN-07-0200.

50. Hoekstra ME, Bornes L, Dijkgraaf FE, Philips D, Pardieck IN, Toebes M, et al. Long-distance modulation of bystander tumor cells by CD8(+) T cell-secreted IFNgamma. Nat Cancer 2020;1(3):291–301 doi 10.1038/s43018-020-0036-4.

51. Gao J, Shi LZ, Zhao H, Chen J, Xiong L, He Q, et al. Loss of IFN-gamma Pathway Genes in Tumor Cells as a Mechanism of Resistance to Anti-CTLA-4 Therapy. Cell 2016;167(2):397–404 e9 doi 10.1016/j.cell.2016.08.069.

52. Shao DD, Xue W, Krall EB, Bhutkar A, Piccioni F, Wang X, et al. KRAS and YAP1 converge to regulate EMT and tumor survival. Cell 2014;158(1):171–84 doi 10.1016/j.cell.2014.06.004.

53. Zhu C, Kim K, Wang X, Bartolome A, Salomao M, Dongiovanni P, et al. Hepatocyte Notch activation induces liver fibrosis in nonalcoholic steatohepatitis. Science translational medicine 2018;10(468) doi 10.1126/scitranslmed.aat0344.

54. Susaki EA, Tainaka K, Perrin D, Yukinaga H, Kuno A, Ueda HR. Advanced CUBIC protocols for whole-brain and whole-body clearing and imaging. Nature protocols 2015;10(11):1709–27 doi 10.1038/nprot.2015.085.

55. Bolger AM, Lohse M, Usadel B. Trimmomatic: a flexible trimmer for Illumina sequence data. Bioinformatics 2014;30(15):2114–20 doi 10.1093/bioinformatics/btu170.

56. Dobin A, Davis CA, Schlesinger F, Drenkow J, Zaleski C, Jha S, et al. STAR: ultrafast universal RNA-seq aligner. Bioinformatics 2013;29(1):15–21 doi 10.1093/bioinformatics/bts635.

57. Anders S, Pyl PT, Huber W. HTSeq--a Python framework to work with high-throughput sequencing data. Bioinformatics 2015;31(2):166–9 doi 10.1093/bioinformatics/btu638.

58. Love MI, Huber W, Anders S. Moderated estimation of fold change and dispersion for RNA-seq data with DESeq2. Genome Biol 2014;15(12):550 doi 10.1186/s13059-014-0550-8.

59. Chen EY, Tan CM, Kou Y, Duan Q, Wang Z, Meirelles GV, et al. Enrichr: interactive and collaborative HTML5 gene list enrichment analysis tool. BMC Bioinformatics 2013;14:128 doi 10.1186/1471-2105-14-128.

60. Foroutan M, Bhuva DD, Lyu R, Horan K, Cursons J, Davis MJ. Single sample scoring of molecular phenotypes. BMC Bioinformatics 2018;19(1):404 doi 10.1186/s12859-018-2435-4.

61. Binder JX, Pletscher-Frankild S, Tsafou K, Stolte C, O’Donoghue SI, Schneider R, et al. COMPARTMENTS: unification and visualization of protein subcellular localization evidence. Database (Oxford) 2014;2014:bau012 doi 10.1093/database/bau012.

62. Cancer Genome Atlas Research Network. Electronic address wbe, Cancer Genome Atlas Research N. Comprehensive and Integrative Genomic Characterization of Hepatocellular Carcinoma. Cell 2017;169(7):1327–41 e23 doi 10.1016/j.cell.2017.05.046.

63. Colaprico A, Silva TC, Olsen C, Garofano L, Cava C, Garolini D, et al. TCGAbiolinks: an R/Bioconductor package for integrative analysis of TCGA data. Nucleic Acids Res 2016;44(8):e71 doi 10.1093/nar/gkv1507.

64. Hanzelmann S, Castelo R, Guinney J. GSVA: gene set variation analysis for microarray and RNA-seq data. BMC Bioinformatics 2013;14:7 doi 10.1186/1471-2105-14-7.

